# Fast and compact matching statistics analytics

**DOI:** 10.1101/2021.10.05.463202

**Authors:** Fabio Cunial, Olgert Denas, Djamal Belazzougui

## Abstract

**Motivation:** Fast, lightweight methods for comparing the sequence of ever larger assembled genomes from ever growing databases are increasingly needed in the era of accurate long reads and pan-genome initiatives. Matching statistics is a popular method for computing whole-genome phylogenies and for detecting structural rearrangements between two genomes, since it is amenable to fast implementations that require a minimal setup of data structures. However, current implementations use a single core, take too much memory to represent the result, and do not provide efficient ways to analyze the output in order to explore local similarities between the sequences.

**Results:** We develop practical tools for computing matching statistics between large-scale strings, and for analyzing its values, faster and using less memory than the state of the art. Specifically, we design a parallel algorithm for shared-memory machines that computes matching statistics 30 times faster with 48 cores in the cases that are most difficult to parallelize. We design a lossy compression scheme that shrinks the matching statistics array to a bitvector that takes from 0.8 to 0.2 bits per character, depending on the dataset and on the value of a threshold, and that achieves 0.04 bits per character in some variants. And we provide efficient implementations of range-maximum and range-sum queries that take a few tens of milliseconds while operating on our compact representations, and that allow computing key local statistics about the similarity between two strings. Our toolkit makes construction, storage, and analysis of matching statistics arrays practical for multiple pairs of the largest genomes available today, possibly enabling new applications in comparative genomics.

**Availability ad implementation:** Our C/C++ code is available at https://github.com/odenas/indexed_ms under GPL-3.0.

## 1 Introduction

Several large-scale projects are under way to assemble the genome of hundreds of new species, and comparing such genomes is crucial for understanding the genetic basis and origin of complex traits and related diseases (for a small sampler, see e.g. [39, 34, 14, 37, 16, 44, 26, 25]). Efficient tools for comparing genome-scale sequences are thus becoming increasingly necessary. The *matching statistics* of a string *S*, called the *query*, with respect to another string *T*, called the *text*, is an array `MS`_*S,T*_ [0..|*S*| − 1] such that `MS`_*S,T*_ [*i*] is the length of the longest prefix of *S*[*i*.. |*S*| − 1] that occurs anywhere in *T* without errors. Since the match can occur anywhere in *T*, matching statistics is robust to largescale rearrangements and horizontal transfers that are common in genomes, and the average matching statistics length over the whole sequence has been used for building consistent whole-genome phylogenies without alignment – and, unlike *k*-mer methods, without parameters [8, 43]. The effectiveness of matching statistics in alignment-free phylogenetics has even motivated variants that allow for a user-specified number of mismatches (for a small sampler see e.g. [2, 27, 30, 40, 41]); and other, seemingly different, distances can be expressed in terms of matching statistics as well [13, 42, 7]. For genomes from the same or from closely related species, matching statistics has been used for computing estimators of the number of substitutions per site, of the number of pairwise mismatches, and of the occurrence of recombination events (see e.g. [23, 21, 19, 20, 10]); finally, the related notion of *shortest unique substring*, defined on a single sequence, has been employed for computing measures of genome repetitiveness [24, 22], and it could be used as a parameter-free method for detecting segmental duplications [31]. Since every position of the query *S* is assigned a match length, matching statistics can reveal ranges of locally high similarity (i.e. of large average matching statistics in the range) induced e.g. by horizontal gene transfer, or conversely ranges of locally low similarity induced by chromosomes of *S* missing from *T*, or by horizontal transfer events that affected *S* but not *T* [24, 23, 20, 11, 12]. See Figures 6 and 7 in the supplement for a concrete example. This idea has been recently applied to targeted Nanopore sequencing, using online matching statistics to eject from the pore a long DNA molecule that is not likely to belong to the species of interest, after having read just a short segment of the molecule [1].

Computing `MS`_*S,T*_ is a classical problem in string processing, and in practice it involves building an index on a fixed *T* to answer a large number of queries *S*. Thus, solutions typically differ on the index they use, which can be the textbook suffix tree, the compressed suffix tree [29] or compressed suffix array, the colored longest common prefix array [17], a Burrows-Wheeler index combined with the suffix tree topology [3, 4], or the *r*-index combined with balanced grammars [6]. In the frequent case where *T* consists of one genome (or proteome), or of the concatenation of few similar genomes or of many dissimilar genomes, the Burrows-Wheeler transform of *T* does not compress well, and the best space-time trade-offs are achieved by the implementation in [4] (see [6] for a runtime comparison, and see Figure 2 in the supplement for a memory comparison). In this paper we develop several practical tools for computing the matching statistics array between genome-scale strings, and for analyzing its values, faster and using less memory than the state of the art.

Specifically, we design a practical variant of the algorithm by [4] that computes MS in parallel on a shared-memory machine, and that achieves approximately a 41-fold speedup of the core procedures and a 30-fold speedup of the entire program with 48 cores on the instances that are most difficult to parallelize. Our implementation takes around 12 minutes to compute the MS between the *Homo sapiens* and the *Pan troglodytes* genomes on a standard 48-core server. We also describe a theoretical variant with better asymptotic complexity, which takes *O*(|*S*| log σ*/t* + (log |*T*| log σ)(log *t* + log log *t* log log |*T*|)) time and 2 |*T*| log σ + *O*(*n*) bits of space when executed on *t* processors, where σ is the integer alphabet of *S* and *T*. To the best of our knowledge, no algorithm for computing matching statistics in parallel existed before.

Then, we implement fast range queries for computing the average and maximum matching statistic value inside a substring of *S*, taking advantage of the compact encoding of `MS`_*S,T*_ introduced by [3]: this encoding takes just 2 |*S*| bits, and allows one to retrieve `MS`[*i*] in constant time for any *i* using just *o*(|*S*|) more bits. In some cases this bitvector is compressible, so our code can operate both on the plain encoding and on its compressed versions. Overall, we can answer queries over arbitrary ranges of the human genome in a few tens of milliseconds, taking just a few extra megabytes of space. No tool for fast range queries over a compact matching statistics encoding existed before.

Finally, we describe a lossy compression scheme that can reduce the size of our compact encoding to much less than 2|*S*| bits when *S* and *T* are dissimilar, by replacing small matching statistics values (that typically arise from random matches) with other, suitably chosen small values. In practice this is most useful in applications that need the matching statistics array of every pair of genomes in a large dataset. The threshold of our lossy compression can be set according to some expected length of matches (see e.g. [20, 21, 23]), or it could be learnt from the distribution of match lengths itself, which usually peaks at noisy values (see e.g. Figure 4 in the supplement). Depending on the threshold, our scheme can shrink the encoding from 40% to 10% of its original size of 2 bits per character, and one of our variants achieves 2% for large thresholds. Another popular data structure in string indexing, the *permuted longest common prefix array* [35], has a similar bitvector encoding and shrinks at similar rates under our scheme in practice.

Our compression method bears some similarities to the *lossless* algorithm by [5], which builds an approximation of the select function on arbitrary bitvectors, and stores corrections: in our case, discarding the corrections would amount to replacing every matching statistics value (regardless of whether it is small or large) with another value (which could be either bigger or smaller) within a user-specified error. This might be undesirable for matching statistics, since there is often an expected length of random matches, and large values that carry information should better be kept intact for downstream analysis. The two lossy schemes are incomparable. In practice the one by [5] tends to produce smaller files, since it has more degrees of freedom; our methods manage to achieve compression rates of similar magnitude in several cases (see Figure 15 in the supplement). The lossless version of [5] *expands* our bitvectors for all settings (see Figure 3 in the supplement).

## 2 Preliminaries and notation

### 2.1 Strings and string indexes

Let Σ = [1..σ] be an integer alphabet, and let *T* ∈ Σ^+^ be a string. We call the *reverse of T* the string 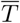 obtained by reading *T* from right to left, and we denote by *f*_*T*_(*W*) the number of occurrences (or frequency) of string *W* in *T*. For reasons of space we assume the reader to be already familiar with the notion of *suffix tree* `ST`_*T*_ = (*V, E*) of *T*, which we do not define here. We just recall that every edge in *E* is labelled with a string of length possibly greater than one, and that a substring *W* of *T* can be extended to the right with at least two distinct characters iff *W* = *ℓ*(*v*) for some internal node *v* of the suffix tree, where *ℓ*(*v*) is the string label of node *v* ∈ *V* obtained by concatenating the label of every edge in the path from the root to *v*. It is well-known that all the nodes in a suffix tree path have distinct frequencies, which decrease from top to bottom. If *u* is a node of the suffix tree of *T*, we use *f*_*T*_(*u*) as a shorthand for *ℓ*_*T*_(*ℓ*(*u*)). We assume the reader to be familiar with the notion of *suffix link* connecting a node *v* with *ℓ*(*v*) = *aW* for some *a* ∈ [1..σ], to a node *w* with *ℓ*(*w*) = *W*. Here we just recall that inverting the direction of all suffix links yields the so-called *explicit Weiner links*. Given an internal node *v* of `ST`_*T*_ and a symbol *a* ∈ [1..σ], it might happen that string *aℓ*(*v*) occurs in *T*, but that it is not the label of any internal node: all such left extensions of internal nodes that end in the middle of an edge or in a leaf are called *implicit Weiner links*. An internal node of `ST`_*T*_ can have several outgoing Weiner links, and every one of them is labelled with a distinct character.

We call *suffix tree topology* a data structure that supports operations on the shape of `ST`_*T*_, like `parent` (*v*), which returns the parent of a node *v*; lca(*u, v*), which returns the lowest common ancestor of nodes *u* and *v*; `leftmostLeaf`(*v*) and `rightmostLeaf` (*v*), which compute the identifier of the leftmost (respectively, rightmost) leaf in the subtree rooted at node *v*; `selectLeaf` (*i*), which returns the identifier of the *i*-th leaf in preorder traversal; `leafRank` (*v*), which computes the number of leaves that occur before leaf *v* in preorder traversal. It is known that the topology of an ordered tree with *n* nodes can be represented using 2*n* + *o*(*n*) bits as a sequence of 2*n* balanced parentheses, and that 2*n* + *o*(*n*) more bits suffice to support every operation described above in constant time [28, 36]. We assume the reader to be familiar also with the Burrows-Wheeler transform of *T* (denoted `BWT`_*T*_ in what follows). Here we just recall that every suffix tree node corresponds to a compact lexicographic interval in the BWT, and that following a Weiner link in the suffix tree, i.e. extending a string *W* = *ℓ*(*v*) to the left with one character, corresponds to the well-known *backward step* from the BWT interval of *W*. We also mention the classical operations `rank`(*T, a, i*), which returns the number of occurrences of character *a* in string *T* up to position *i*, inclusive; and `select`(*T, a, i*), which returns the position of the *i*-th occurrence of *a* in *T*.

In what follows we omit subscripts that are clear from the context, and we use 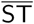 and 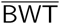 as shorthands for 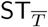 and 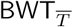, respectively.

### 2.2 Matching statistics in small space

As mentioned, given a query string *S* ∈ Σ^*m*^, we call *matching statistics* `MS`_*S,T*_ [0..*m* − 1] an array such that `MS`_*S,T*_ [*i*] is the length of the longest prefix of *S*[*i*..*m*−1] that occurs somewhere in *T* without errors. In this paper we work with the compact representation of `MS`_*S,T*_ as a bitvector `ms`_*S,T*_ of 2|*S*| bits, which is built by appending, for each *i* ∈ [0..|*S*| − 1] in increasing order, `MS`_*S,T*_ [*i*] − `MS`_*S,T*_ [*i* − 1] + 1 zeros followed by a one [3]. `MS`_*S,T*_ [−1] is assumed to be one. Since the number of zeros before the *i*-th one in ms equals *i* + `MS`[*i*], one can compute `MS`[*i*] for any *i*∈ [0.. |*S*| −1] using select operations on ms. We also work with the algorithm by [3], which we summarize here for completeness. This offline algorithm computes `ms` using both a backward and a forward scan over *S*, and it needs in each scan just `BWT` with rank support, and the topology of `ST`, or just 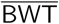 with rank support, and the topology of 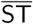. The two phases are connected via a bitvector `runs` [1..|*T*| −1], such that `runs` [*i*] = 1 iff `MS`[*i*] = `MS`[*i* − 1] − 1, i.e. iff there is no zero between the *i*-th and the (*i* − 1)-th ones in `ms`.

First, we scan *S* from right to left, using `BWT` with rank support, and the suffix tree topology of *T*, to determine the runs of consecutive ones in `ms`. Assume that we know the interval [*i*..*j*] in `BWT` that corresponds to substring *W* = *S*[*k*..*k* + `MS`[*k*] −1], as well as the identifier of the proper locus *v* of *W* in the topology of `ST`. We try to perform a backward step using character *a* = *S*[*k* −1]: if the resulting interval [*i*′..*j*′] is nonempty, we set `runs` [*k*] = 1 and we reset [*i*..*j*] to [*i*′..*j*′]. Otherwise, we set `runs` [*k*] = 0, we update the BWT interval to the interval of `parent` (*v*) using the topology, and we try another backward step with character *a*.

In the second phase we scan *S* from left to right, and we build`ms` using 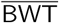 with rank support, the suffix tree topology of 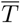, and bitvector`runs`. Assume that we know the interval [*i*..*j*] in 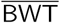 that corresponds to substring *W* = *S*[*k*..*h* −1] such that `MS`[*k* −1] = *h* −*k* but `MS`[*k*] ≥ *h* −*k*. We try to perform a backward step with character *S*[*h*]: if the backward step succeeds, we continue issuing backward steps with the following characters of *S*, until we reach a position *h*^*^ in *S* such that a backward step with character *S*[*h*^*^] from the interval [*i*^*^..*j*^*^] of substring *W* ^*^ = *S*[*k*..*h*^*^ −1] in 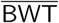 fails. At this point we know that `MS`[*k*] = *h*^*^ −*k*, so we append *h*^*^−*k*−`MS`[*k*−1] + 1 = *h*^*^−*h* + 1 zeros and a one to ms. Moreover, we iteratively reset the current interval in 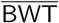 to the interval of `parent` (*v*^*^), where *v*^*^ is the proper locus of *W* ^∗^ in 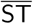, and we try another backward step with character *S*[*h*^*^], until we reach an interval [*i*′..*j*′] for which the backward step succeeds. Let this interval correspond to substring *W* ′ = *S*[*k*′..*h*^*^− 1]. Note that `MS`[*k*′] > `MS`[*k*′ −1] 1 and `MS`[*x*] = `MS`[*x* −1] −1 for all *x* ∈ [*k* + 1..*k*′ −1], so *k*′ is the position of the first zero to the right of position *k* in`runs`, and we can append *k*′ − *k* − 1 ones to`ms`. Finally, we repeat the whole process from substring *S*[*k*′..*h*^*^] and its interval in 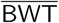.

## 3 Computing MS in parallel

It is natural to try and parallelize the construction of MS when query strings are long. In the case of proteomes, or of concatenations of several small genomes, reads, or contigs, one could just split the query in chunks of approximately equal size along concatenation boundaries, and process each chunk in parallel. For the large, contiguous genome assemblies that are increasingly achievable with long reads, one could compute MS in parallel for each chromosome, but chromosomes might have widely different lengths and their number might be much smaller than the number of cores available. In this section we describe algorithms for computing MS in parallel for long query strings, without assuming that they are the concatenation of shorter strings.

We work in the concurrent read, exclusive write (CREW) model of a parallel random access machine, in which multiple processors are allowed to read from the same memory location at the same time, but only one processor is allowed to write to a memory location at any given time. Let *S*^0^, …, *S*^*t*−1^ be a partition of *S* into *t* blocks of equal size, and let *p*_*i*_ be the first position of block *S*^*i*^. Once *S* and `runs` are loaded in memory, one can build `ms` in parallel with *t* threads, by computing `MS`[*p*_*i*_, …, *p*_*i*+1_− 1] independently for each *i*: this works since every thread can safely read the suffix of *S* to the right of its own block, as well as the corresponding positions of `runs.` Recall however that the algorithm outputs a compact representation of array `MS`, rather than array `MS` itself. Let `ms`^*i*^ be the bitvector representation of `MS`[*p*_*i*_, …, *p*_*i*+1_ − 1]. Thread *i* computes `ms`^*i*^ starting from position *p*_*i*_ of *S*, and it appends to the beginning of `ms`^*i*^ a sequence of `MS`[*p*_*i*_] zeros and a one; however, in the final bitvector `ms`, such a sequence of bits should be replaced by a sequence of `MS`[*p*_*i*_] −`MS`[*p*_*i*_ −1] + 1 zeros and a one. We perform this correction in a final pass, in which a single thread concatenates all output bitvectors. Specifically, we use `MS`[|*S*^0^| − 1] = | `ms`^0^| −2 |*S*^0^| + 1 to correct the first run of zeros of `ms`^1^, and so on for the other blocks.

We compute bitvector `runs` with *t* parallel threads, as follows. Let *R*^0^, …, *R*^*t*−1^ be the partition of `runs` induced by blocks *S*^0^, …, *S*^*t*−1^. Thread *i* executes the algorithm for computing `runs` independently just inside blocks *S*^*i*^ and *R*^*i*^, starting with filling the last bit of *R*^*i*^. Assume that thread *i*, while proceeding from right to left, sets bit *b*_*i*_ of *R*^*i*^ to zero, and that it sets *R*^*i*^[*b*_*i*_ + 1.. |*R*^*i*^ | −1] to all ones. All the bits that thread *i* sets in *R*^*i*^[0..*b*_*i*_] are correct, since they can be decided without looking at blocks *S*^*i*+1^, …, *S*^*t*−1^. However, to decide the value of bits *R*^*i*^[*b*_*i*_ + 1.. |*R*^*i*^| −1] one needs to look at the blocks that follow *S*^*i*^. We call *marked* the last block *R*^*t*−1^, as well as any block *R*^*i*^ that contains a zero after this phase. If *R*^*i*^ is marked, let *W*^*i*^ = *S*[*p*_*i*_..*p*_*i*_ + `MS`[*p*_*i*_] −1]: then, thread *i* stores the BWT interval and the topology identifier of the locus of *W*_*i*_ in `ST`_*T*_.

In practice we expect *b*_*i*_ to be close to |*R*^*i*^| − 1, and we expect most blocks to be marked. However, there could be an *R*^*i*^ that contains no zero after this phase. Thus, we have to run a second phase in which, for every marked block *R*^*i*^, we start a thread that updates all the one-bits, in all blocks between *R*^*i*−1^ and the rightmost marked block *R*^*j*^ before *R*^*i*^, including the suffix of *R*^*j*^ after its last zero. We perform this correction using the information stored in the previous phase. Note that this strategy might result both in using fewer than *t* threads (since we issue just one thread per marked block), and in linear time per thread, since the number of one-bits that a thread might have to update could be proportional to |*S*|.

These problems occur when *S* and *T* have long exact matches, and they become extreme when *S* = *T*. Thus, we experiment with the pairs of similar genomes and proteomes described in Section 1 of the supplement. The construction of both `runs` and `ms` scales well on genomes and proteomes of similar species, although achieving the ideal speedup gets more difficult as the number of threads increases (Figure 2). Correcting the `runs` bitvector takes a negligible fraction of the total time for processing `runs`, even for similar genomes, and it takes more time for proteomes than for genomes, probably because the proteomes of related species are more similar to one another than their genomes (see Figure 1 in the supplement). The fraction of time spent in correcting `runs` grows with the number of threads, probably because more threads imply shorter blocks, and shorter blocks are more likely to intersect with exact matches between *S* and *T*, or to be fully contained in them. Correcting the `ms` bitvector takes even less time than correcting `runs`. Even running the algorithm on the very similar pairs of chromosome 1 from different human individuals shows the same trends (Figure 1 in the supplement). When *S* = *T*, correcting `runs` uses just one thread, since only the last block is marked, and it takes time proportional to |*S*| (*t* −1)*/t* (Figure 2); building `ms` uses all threads, but each one of them has to process the whole suffix of *S* that starts from its block, thus there is no speedup with respect to the sequential version. In the following section we describe a way to achieve better asymptotic complexity even when *S* = *T*.

**Figure 1:**
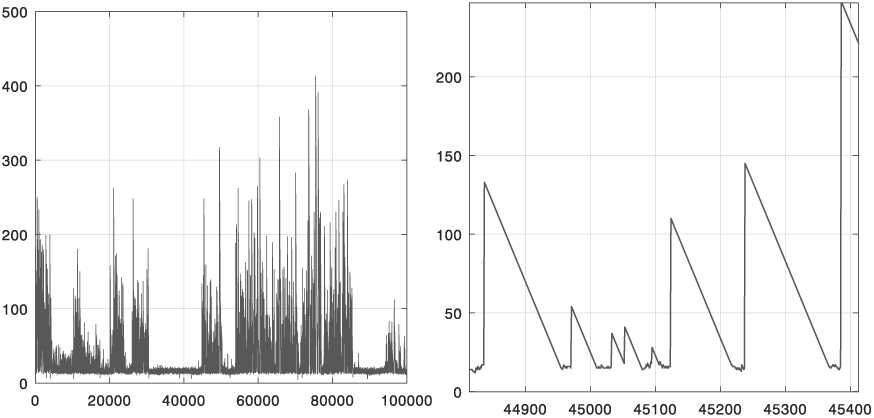
Values of matching statistics (vertical axis) in a range of positions along human chromosome 1 (horizontal axis). Query: *Homo sapiens*. Text: *Pan troglodytes*. The right panel is a zoom-in of the left panel. See Figure 6 in the supplement for a bigger range. The PLCP array of a single genome (defined in Section 4) has a similar shape.

**Figure 2:**
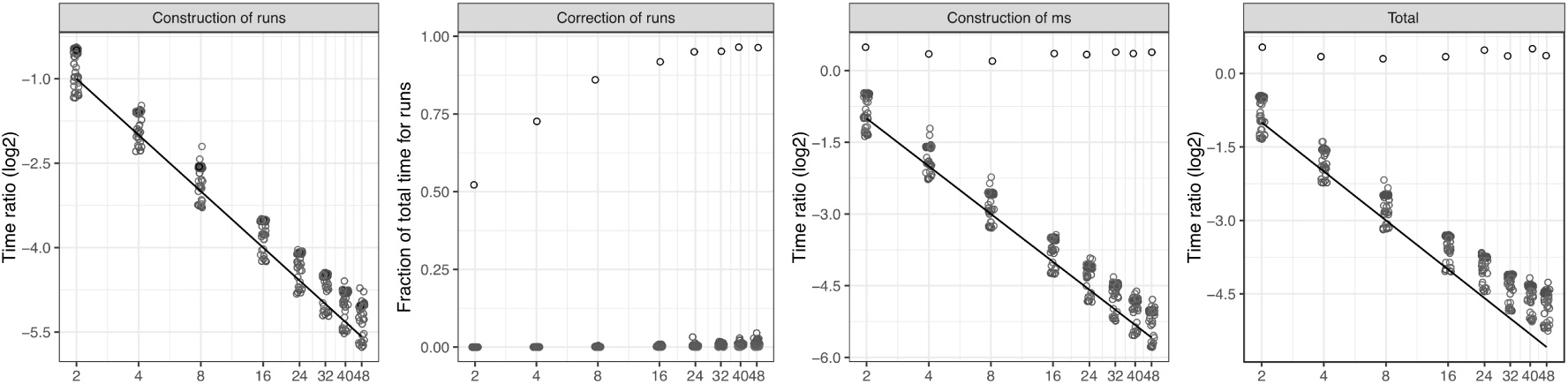
Scaling of our parallel implementation as the number of threads *t* increases. Line: ideal scaling 1*/t*. Circles along the line: genomes of similar species, proteomes of similar species, pairs of human chromosome 1 from distinct random individuals. Circles far from the line: identical query and text (human chromosome 1 from two random individuals). Vertical axis: time of the parallel implementation divided by the time of the sequential implementation. Correction of `ms` is not shown since it is negligible. See Figure 1 in the supplement for more details.

### 3.1 Better asymptotic complexity

Another way of computing `runs` in parallel could be by performing a backward search from the end of every block *S*^*i*^ using 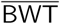, by mapping the resulting interval to the corresponding interval in `BWT`, and by starting the computation of each block from such intervals. This naive approach has the disadvantage of requiring linear time per thread in the worst case to compute the initial BWT intervals of the blocks, and of needing to translate intervals from one BWT to the other. However, the general idea can be used to achieve better complexity, as follows:

#### Lemma 1.

*Let T be a string on alphabet* [1..σ], *and assume that we have a representation of* `SA`_*T*_ *that support suffix array and inverse suffix array queries in O*(*p*) *time, and a representation of* `ST`_*T*_ *that support* `weinerLink` *and* `parent` *queries in O*(*q*) *time. Given a query string S and t processors, we can compute* `ms` _*S,T*_ *in time O*(|*S*| · *q/t* + log log |*T*|*· p* log *t*) *using O*(|*S*| + *t* log |*T*|) *bits of working space*.

*Proof*. Without loss of generality, we assume that *t* is a power of two. To compute the `runs` bitvector, we proceed as follows. First, we split *S* into *t* blocks *S*^0^, …, *S*^*t*−1^, and we build the BWT interval of every block *S*^*i*^ in parallel, spending *O*(|*S*|*q/t*) time overall. Then, we build the BWT interval of every disjoint *superblock* that consists of 2^*j*^ adjacent blocks, for all *j*∈ [0.. log *t* − 1], in log *t* phases. In phase *j* we compute the BWT interval of every one of the *t/*2^*j*^ superblocks in parallel, by merging the BWT intervals of the two smaller superblocks from the previous phase that compose it. Every such merge can be performed in *O*(*p* log log |*T*|) time using a data structure that takes *O*(|*S*|) additional bits of space [15], so all merging phases take *O*(log *t p* log · log |*T*|) time in total. Note that storing the BWT intervals of all superblocks from all phases takes just *O*(*t* log |*T*|) bits of working space. Then we compute, for every *i*∈ [0..*t*−1], the largest *j* ≥*i* such that *S*^*i*+1^ *S*^*j*^ occurs in *T* (we call *g*(*i*) such a value of *j* in what follows). This can be done by assigning a processor to every block *S*^*i*^, and by making the processor merge the BWT intervals of *O*(log *t*) pairs of superblocks computed previously. This takes again *O*(log *t · p* log log |*T*|) time overall. Finally, for every *S*^*i*^ in parallel, we try to extend the BWT interval of *S*^*i*+1^ *S*^*g*(*i*)^ inside the next block *S*^*g*(*i*)+1^, by performing *O*(*S /t*) backward steps in overall *O*(*S q/t*) time.

We use the resulting intervals for computing the block of `runs` that corresponds to every block *S*^*i*^, independently and in parallel, in overall *O*(|*S*|*q/t*) time. To compute the block of ms that corresponds to each *S*^*i*^, independently and in parallel, we first need to compute the interval in 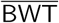 of the longest prefix of *S*^*i*^, …, *S*^*t*−1^ that occurs in *T* : we compute all such intervals using the same superblock approach described above.

Plugging into Lemma 1 some well-known suffix array representations, we can get *O*(|*S*| log σ*/t* + log log |*T*| log |*T*| log σ log *t*) time and 2 |*T*| log σ + *O*(*n*) bits of space for an integer alphabet of size σ, or *O*(|*S*|*/t* + log log |*T*| log *t*) time and *O*(|*T*| log |*T*|) bits of space for an alphabet that is polynomial in |*T*|.

We can further improve on the complexity of every step of Lemma 1, by using a parallel rather than a sequential algorithm for computing the BWT interval of *V W*, given the BWT intervals of *V* and of *W*. Specifically, we use the algorithm by [15], which runs in *O*(*p* log_*t*_ log |*T*|) time with *t* processors, taking again *O*(|*S*|) bits of working space:

#### Lemma 2.

*Given the assumptions of Lemma 1, we can compute* `ms`_*S,T*_ *in time O*(|*S*|*· q/t* + log log |*T*|*· p* log log *t* + *p* log *t*) *using O*(|*S*| + *t* log |*T*|) *bits of working space*.

*Proof*. To build the BWT interval of every superblock, we proceed as follows. In phase *j* we have to merge *t/*2^*j*^ pairs of BWT intervals (one for each superblock of 2^*j*^ blocks), thus we can afford to allocate 2^*j*^ processors to each merge: it is easy to see that this yields *O*(log log |*T*|*· p* log log *t* + *p* log *t*) time overall.

Then, to compute *g*(*i*) for each *i*, we proceed as follows. If we had to solve the problem just for the blocks whose ID is a multiple of 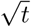, we could allocate 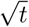 processors to each task and be done in 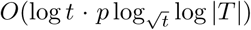 time, which is *O*(*p* log log *T*). More generally, we could organize the computation in *O*(log log *t*) iterations: at iteration *j* we solve the problem for all remaining blocks whose ID is a multiple of 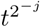, for increasing *j* (i.e. from larger to smaller offsets). Thus, every block *i* that we want to solve at iteration *j* lies between two blocks *i*_*x*_ < *i* < *i*_*x*+1_ that we solved at iteration *j*− 1. Clearly there are 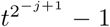 total blocks between 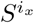 and 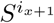 (excluded), thus in the current iteration we have to compute the solution for 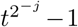 blocks that lie between 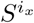 and 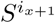. Moreover, since we are dealing with matching statistics, *g*(*i*_*x*_) ≤ *g*(*i*) ≤ *g*(*i*_*x*+1_), and the sum of *g*(*i*_*x*+1_) − *g*(*i*_*x*_) over all *x* is at most *t*. It follows that, if we assigned *g*(*i*_*x*+1_) −*g*(*i*_*x*_) processors to compute each solution between 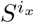 and 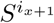, we would end up using 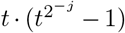 processors in total: since we have just *t* processors, we should thus assign 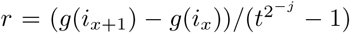 processors to each solution^1^. Since *g*(*i*_*x*_) ≤ *g*(*i*) ≤ *g*(*i*_*x*+1_), we need to merge just *O*(log(*g*(*i*_*x*+1_) −*g*(*i*_*x*_))) pairs of superblocks to compute the solution for any *S*^*i*^, thus the total running time of one iteration is *O*(log(*g*(*i*_*x*+1_) − *g*(*i*_*x*_)) · *p* log_*r*_ log |*T*|) using the parallel algorithm by [15]: it is easy to see that this is *O*(*p* log log |*T*|), thus we get the claimed bound over *O*(log log *t*) iterations. □

By plugging into Lemma 2 the same data structures as before, we can get *O*(|*S*| log σ*/t* +log log |*T*| log |*T*| log σ log log *t* + log |*T*| log σ log *t*) time and 2 |*T*| log σ + *O*(*n*) bits of space, or *O*(|*S*|*/t* + log log |*T*| log log *t* + log *t*) time and *O*(|*T*| log |*T*|) bits of space, which is comparable to the complexity of prefix matching queries described by [15].

## 4 Compressing the MS bitvector

Even though `ms` _*S,T*_ takes just 2|*S*| bits, storing the bitvector of every pair of genomes in a large dataset for later analysis and querying might still require too much space overall. Real `ms` bitvectors, however, have several features that could be exploited for lossless compression. Specifically, if *S* and *T* are similar, they are likely to contain long *maximal exact matches* (MEMs), i.e. triplets (*i, j, f*) such that *S*[*i*..*i*+ *ℓ*−1] = *T* [*j*..*j* + *ℓ*−1], *S*[*i*−1] ≠ *T* [*j* −1] and *S*[*i*+ *ℓ*] ≠ *T* [*j* + *ℓ*]. In practice, MEMs tend to be surrounded in *S* by regions with short matches with *T* (see the example in Figure 1), so `MS`[*i* − 1] is likely to be short, and the run of *ℓ* − `MS`[*i* − 1] + 1 zeros induced by *S*[*i*..*i* + *ℓ* − 1] in `ms` is likely to be long; moreover, in practice `MS`[*i*′] = `MS`[*i*′ − 1] for all *i*^*t*^ ∈ [*i* + 1..*i* + *f* − *k*] for some small *k* (see again Figure 1). Thus, every MEM is likely to induce a long run of zeros followed by a long run of ones in `ms`, and if *S* and *T* share several long MEMs, run-length encoding `ms` might save space. Another property of real `ms` bitvectors is that the length of a long run of zeros tends to be similar to the length of the following long run of ones, since *ℓ* − `MS`[*i* − 1] + 1 − *ℓ* + *k* = *k* − `MS`[*i* − 1] + 1 is likely to be small (see Figures 18 and 19 in the supplement). So, given a pair (*z*_*i*_, *o*_*i*_) representing a run of *z*_*i*_ zeros and the following run of *o*_*i*_ ones, and given an encoder *δ*, one might encode the pair as *δ*(*z*_*i*_)*δ*(*o*_*i*_ − *z*_*i*_) if *z*_*i*_ is large, and as *δ*(*z*_*i*_)*δ*(*o*_*i*_) otherwise.

Run-length encoding the bitvectors of pairs of genomes from human individuals using e.g. the `RLEVector` data structure by [38] yields compression rates of about 20 (Figure 3, right panel), and compressing the same bitvectors with the `rrr_vector` data structure from the SDSL library [18] (which implements the RRR scheme by [33]) yields compression rates of about 6 (see Figure 8 in the supplement). However, the bitvectors of *pairs of genomes from different species* are recalcitrant to compression, even when the species are related: runlength encoding *expands* those files by a factor of two (Figure 3, insert in the left panel), and RRR expands most of them slightly (by a factor of 1.1), and manages to compress just few pairs with rate 1.25 (Figure 8 in the supplement). The same happens with pairs of artificial strings with controlled mutation rate (see Figures 16, 17 in the supplement).

**Figure 3:**
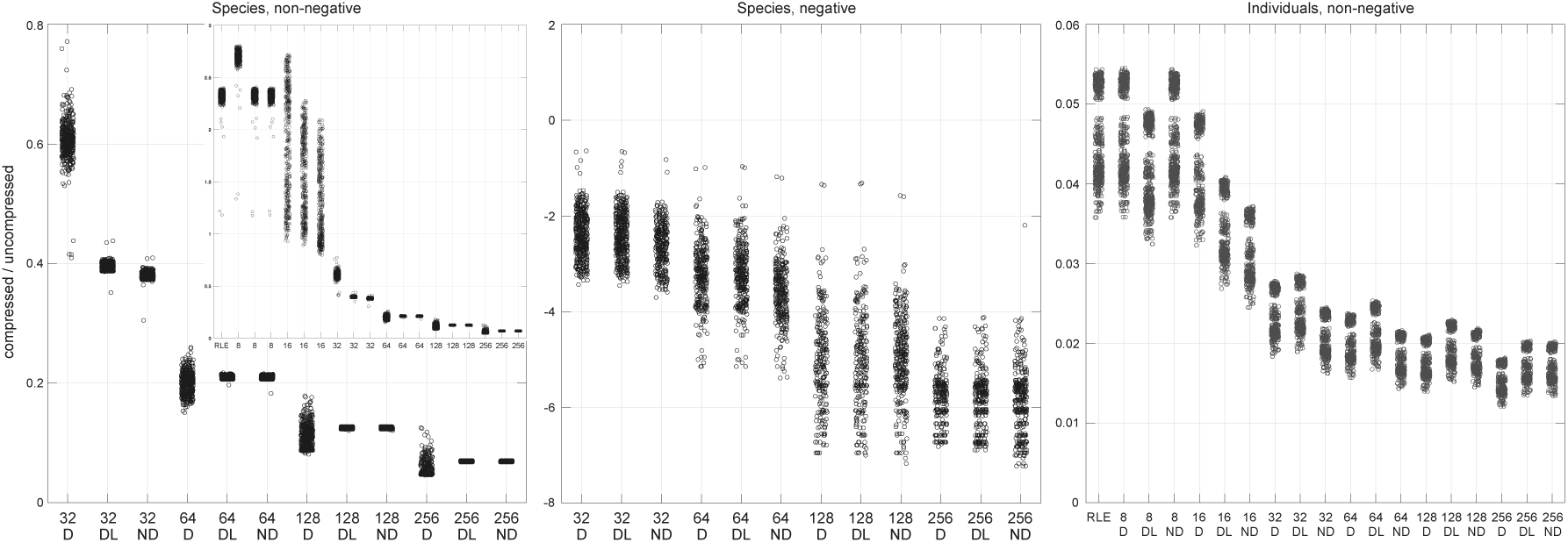
Ratio between the size of the `RLEVector` data structure by [38] built on a permuted `ms` `bit_vector`, and the size of the `bit_vector` data structure from SDSL built on the original `ms` bitvector, for the D, DL, and ND lossy variants on pairs of genomes from different species and on pairs of genomes from human individuals, allowing and disallowing negative MS values. Size is measured on disk. The vertical axis in the middle panel shows negative powers of ten. Computation is exact for windows with up to 300 zeros and 300 ones, then it uses the first greedy strategy described in the text.

In some applications, including genome comparison, short matches are considered noise by the user, and the precise length of a match can be discarded safely as long as we keep track that at that position the match was short. Given an array `MS`_*S,T*_ and a user-defined threshold *τ*, let a *thresholded matching statistics array* `MS`_*S,T,τ*_ be such that `MS`_*S,T,τ*_ [*i*] = `MS`_*S,T*_ [*i*] if `MS`_*S,T*_ [*i*] ≥ *τ*, and `MS`_*S,T,τ*_ [*i*] equals an arbitrary (possibly negative) value smaller than *τ* otherwise^2^. This notion is symmetrical to the one defined by [9], which discards instead long MS values in order to prune the suffix tree topologies and to make the data structures smaller. Given an encoder *δ*, we are interested in the `MS`_*S,T,τ*_ array whose `ms`_*S,T,τ*_ bitvector takes the smallest amount of space when encoded with *δ*. In what follows, we drop *S* and *T* from the subscripts whenever they are clear from the context.

Note that every `ms`_*S,T,τ*_ is a permutation of `ms`_*S,T*_, since the two bitvectors must contain the same number of zeros and ones. Moreover, if `MS`[*x*] ≥ *τ* corresponds to the one-bit at position *y* in ms, then every `ms`_*τ*_ must also have a one at position *y*, which corresponds to `MS`_*τ*_ [*x*] and is preceded by the same number of ones and zeros as position *y* in `ms` (this follows from the fact that `MS`[*x*] = `select` (`ms`, *x*, 1) − 2*x*). Let *x*_0_, …, *x*_*k*−1_ be the sequence of all and only the positions of *S* whose MS value is at least *τ*, and let *y*_0_, …, *y*_*k*−1_ be the sequence of the corresponding one-bits *y*_*i*_ = `select` (`ms`, *x*_*i*_, 1) in `ms`. Clearly it can happen that *x*_*i*+1_ = *x*_*i*_+1; if this does not happen, then `MS`[*x*_*i*_] must be equal to *τ*, and `ms` [*y*_*i*_ + 1] must be a one and `ms` [*y*_*i*+1_ − 1] must be a zero, both in `ms` and in any `ms`_*τ*_. Thus, if we compress `ms`_*τ*_ by deltacoding the length of every run, we can build an`ms` _*τ*_ that is smallest after compression, by concatenating a permutation of every such interval [*y*_*i*_..*y*_*i*+1_ − 1] of `ms` that is smallest after compression, as well as of the non-empty intervals [0..*y*_0_ − 1] and [*y*_*k*−1_ +1..2|*S*|] (and all such permutations can be computed in parallel).

Assume that we want to compute a smallest permutation of window [*y*_*i*_..*y*_*i*+1_ −1], where `MS`[*x*_*i*_] = *τ* and every run is delta-coded in isolation. Clearly we could just replace the window with 1^*p*^0^*q*^, where *p* (respectively, *q*) is total the number of ones (respectively, zeros) in the window; this could make some MS values negative, thus the resulting `ms`_*τ*_ might not be a valid MS bitvector, and before replacing `ms` with `ms` _*τ*_ one should make sure that any implementation that used `ms` handles negative values correctly. Building an MS bitvector without negative values is easy:

### Lemma 3.

*Given an interval* [*y*_*i*_ + 1..*y*_*i*+1_] *of* `ms`, *with z total zeros and o total ones, we can compute a smallest permutation with no negative value in O*(*zoτ* ^2^) *time and words of space*.

*Proof*. Every permutation of the interval can be represented as a sequence of pairs (*z*_0_, *o*_0_), (*z*_1_, *o*_1_), …, (*z*_*k*_, *o*_*k*_) for some *k* ≥ 0, where *z*_*i*_ is the length of a run of zeros, *o*_*i*_ is the length of a run of ones, *z*_0_ ≥ 0, *z*_*i*_ > 0 for all *i >* 0, and *o*_*i*_ > 0 for all *i* ≥ 0. We work with the sequence of *cumulative pairs* (*Z*_0_, *O*_0_), (*Z*_1_, *O*_1_), …, (*Z*_*k*_, *O*_*k*_), where 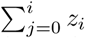 and 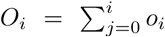. Given a pair (*Z*_*i*_, *O*_*i*_), we use `MS`(*Z*_*i*_, *O*_*i*_) as a shorthand for *τ* +*Z*_*i*_ − *O*_*i*_ (i.e. the MS value that corresponds to the last one-bit of the pair), and we say that the pair is *valid* iff it satisfies *Z*_*i*_ *< z, O*_*i*_ < *o*, and `MS`(*Z*_*i*_, *O*_*i*_) ∈ [0..*τ* − 1]. We draw a directed arc from every valid pair (*Z*_*i*_, *O*_*i*_) to every other valid pair (*Z*_*j*_, *O*_*j*_) such that *Z*_*j*_ > *Z*_*i*_, − *O*_*j*_ > *O*_*i*_, and `MS`(*Z*_*i*_, *O*_*i*_) + *Z*_*j*_ − *Z*_*i*_ − 1 *< τ* (this is the MS value of the first one-bit in the last run of ones in the pair), and we assign cost *δ*(*Z*_*j*_ − *Z*_*i*_) + *δ*(*O*_*j*_ − *O*_*i*_) to the arc. Moreover, we add the invalid pair (*z, o*), we connect it to every valid pair (*Z*_*i*_, *o* − 1), and we assign cost *δ*(*z* − *Z*_*i*_) to the arc. A *start pair* (*Z*_*i*_, *O*_*i*_) is a valid pair with *Z*_*i*_ = 0, and it is assigned cost *δ*(*O*_*i*_). A permutation of smallest size corresponds to a path in the resulting DAG *G* = (*V, E*), from a start pair to pair (*z, o*), that minimizes the sum of the costs of its arcs plus the cost of the start pair. This can be derived by computing, for every node *v* ∈ *V* that does not correspond to a start pair, quantity *f* (*v*) = min {*f* (*u*) + *c*(*u, v*) : (*u, v*) ∈ *E*}, using dynamic programming over the topologicallysorted DAG. □

One can easily modify this construction to enforce MS values in the permuted interval to be at least a positive number, rather than zero. To make compression faster in practice, we fix *z* and *o* to a large value and, for every *τ* used by the target application, we precompute and store a variant of the DAG that answers every possible query of length at most *z* + *o*: in addition to (*z, o*), this variant includes every pair (*Z*_*i*_, *O*_*i*_) with `MS`(*Z*_*i*_, *O*_*i*_) ≥ *τ*, it connects it to all valid pairs (*Z*_*j*_, *O*_*i*_ − 1) as described for (*z, o*), and it computes the min-cost path to every node. To permute a window with *z*′ zeros and *o*′ ones such that *z*′≤ *z*, ≤ *o*′ *o*, and *z*′+*o*′ ≤ *z*+*o*, we go to node (*z*′, *o*′) in the DAG and we backtrack along an optimal precomputed path. If (*z*′, *o*′) does not belong to the DAG, we select a valid in-neighbor (*Z*_*i*_, *O*_*i*_) of (*z*′, *o*′) using a greedy strategy (for example the neighbor that maximizes *g*_*i*_ = *z*′ − *Z*_*i*_ + *o*′ − *O*_*i*_ or *g*_*i*_*/*(*δ*(*z*′ − *Z*_*i*_) + *δ*(*o*′ − *O*_*i*_))): if (*Z*_*i*_, *O*_*i*_) belongs to the DAG, we backtrack, otherwise we take another greedy step. In what follows, we label this approach “ND”.

As mentioned, in real MS bitvectors the length of a run of zeros and of the following run of ones tend to be similar: we can take this into account by setting the cost of an arc between (*Z*_*i*_, *O*_*i*_) and (*Z*_*j*_, *O*_*j*_) to *δ*(*x*)+*δ*(*g*(*y*|*x*)), where *x* = *Z*_*j*_ − *Z*_*i*_, *y* = *O*_*j*_ − *O*_*i*_, and *g*(*y x*) is the following map: since *y* – *x* ≥ 1 − *x*, we map all the negative values of *y* − *x* to the even integers up to 2(*x* − 1) in increasing order of |*y* − *x*|, we map the positive values of *y* − *x* up to *x* − 2 to the odd integers ≥ 3, and we map every remaining value of *y* − *x* to *y*. We use integer one to encode *y* = *x*. Recall that the interval of `ms` that we want to permute is [*y*_*i*_..*y*_*i*+1_ − 1], where *y*_*i*_ belongs to a (possibly long) run of one-bits, and *y*_*i*+1_ is the first one-bit of a (possibly long) run. We might not want to alter the lengths of such runs of ones, so we might be interested in permuting just the subinterval [*p*..*q*] where *p* is the first zero after *y*_*i*_ and *q* is the last one before *y*_*i*+1_ (if negative values of MS are allowed, the trivial scheme of writing all the ones at the beginning of [*p*..*q*] cannot be used, since it would alter the length of the run of *y*_*i*_). We call this variant “D” in what follows. Since in practice the correlation between the length of a run of zeros and the following run of ones is strong only for long runs, one might want to encode a run of *x* zeros ad the following run of *y* ones as *δ*(*x*) + *δ*(*g*(*y x*)) only when *x* ≥ *τ*, and to encode it as *δ*(*x*) + *δ*(*y*) otherwise. This would require permuting just [*p*..*q*], but in an optimal way with respect to the latter encoding. We call this variant “DL” in what follows.

When *S* and *T* are dissimilar, run-length compressing the permuted `ms` bitvectors expands them for small values of *τ* when negative MS values are not allowed (Figure 3, insert in the left panel). For *τ* = 32, run-length encoding most of our `ms`_*τ*_ variants shrinks the bitvector to approximately 40% of its original size, and increasing *τ* progressively brings its size down to 10% of the original. We do not detect any clear difference in performance between the variants, with D being significantly smaller in some but not all cases (Figure 11 in the supplement). A detailed analysis of how the permutation schemes compare when varying the similarity between query and text is provided in Figures 16, 17 in the supplement. For pairs of genomes from human individuals, run-length encoding the original `ms` bitvector already brings its size down to approximately 4.5% of the original, and increasing *τ* shrinks the bitvectors to 2% of the input (Figure 3, right panel). Allowing for negative MS values compresses some pairs of genomes from different species already at *τ* = 16, and for *τ* ≥ 32 it shrinks the bitvector to approximately 2% of the original (Figure 3, center panel). Negative MS values do not give any significant gain for genomes of individuals (data not shown). Pairs of proteomes display similar trends, but this time run-length encoding is able to compress some `ms` bitvectors, and *τ* = 8 is enough to compress most pairs (Figures 9, 10 in the supplement). Finally, we test our lossy compression on the *permuted longest common prefix array* (PLCP) of the genomes in our dataset, since this data structure is amenable to a compact encoding that is very similar to `ms` [35]: we observe again a shrinkage from 40% to 10% of the original size when setting *τ* ≥ 32 (Figure 14 in the supplement).

Clearly, when very few MS values are above threshold, storing just those values might take less space than compressing the ms bitvector. We call *A*_*τ*_ a scheme that stores every MS value at least *τ* and its position in the minimum number of bits necessary to encode the respective numbers, and we call *B*_*τ*_ a scheme in which every MS value at least *τ* is stored in log *M* bits and every position is stored in log *L* bits, where *M* is the maximum observed MS value and *L* is the length of the query. Accessing MS from such structures might be much slower than using the `ms` bitvector with select queries. For *τ* ≥ 32, large values are rare enough in the MS arrays of genomes from different species that most compressed bitvectors take much more space than *A*_*τ*_ or *B*_*τ*_ (Figure 12 in the supplement). When negative values of MS are allowed, however, the permuted bitvectors of several pairs of genomes become smaller than or comparable to *A*_*τ*_ and *B*_*τ*_ (Figure 13 in the supplement), and for pairs of individuals the permuted bitvectors are always two or three orders of magnitude smaller than *A*_*τ*_ and *B*_*τ*_, since 80% or more of all MS values are above threshold for every *τ* (Figure 12 in the supplement).

### 4.1 Compressing frequency and position arrays

Given a position *i* of the query *S*, knowing the frequency of the matching statistics string *S*[*i*..*i*+ `MS`_*S,T*_ [*i*] − 1] in the text *T* can be useful in genome-genome and read-genome comparison, since it can tell for example whether the longest match belongs to an exact repeat of *T* (see e.g. Figure 22 in the supplement). One might also want to keep the exact values of MS just for the positions with low frequency, i.e. one might want to compute a *frequency thresholded* MS array in which every window of ms between two low-frequency one-bits can be permuted to achieve compression. We define the *frequency array* `F`_*S,T*_ [0..*m* − 1] to be such that `F`_*S,T*_ [*i*] = *f*_*T*_ (*S*[*i*..*i* + `MS`_*S,T*_ [*i*] − 1]).

The algorithm for computing ms described in Section 2.2 can be easily adapted to compute the `F` array as well. Specifically, during the first scan of *S* (from right to left), we set runs[*i*] = 0 iff either: (1) `MS`[*i*] ≠ `MS`[*i*−1]−1, as before, or (2) if `MS`[*i*] = `MS`[*i* − 1] − 1 but `F`[*i*] ≠ `F`[*i* −1]. We can detect the latter case since we know the frequency of the current string after every backward step. Consider now the first few operations of the second scan of *S* (from left to right): we managed to match some prefix of *S*, and we are now witnessing a Weiner link fail from some node *v* of 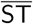, thus we move to the parent *u* of *v*. Node *u* must have a different frequency in *T* than *v*, so we know that the first parent operation we take leads to the position *i* of the first zero-bit from the left in the `runs` bitvector. We can measure *f* (*u*) with the topology, and we know that `F`[*j*] = *f* (*v*) for all *j* ∈ [0..*i* −1]. We repeat this process after every parent operation. At some point the Weiner link succeeds, so we derive the updated frequency from the BWT and we restart the whole process. This algorithm can be parallelized using the same methods as in Section 3.

When *S* and *T* are similar, most matches are long and the values of `F` are more likely to be small or equal to one (and vice versa when *S* and *T* are dissimilar), thus delta-coding `F` achieves better compression when *S* and *T* are similar. Moreover, assume that most edges of 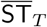 are labelled by long strings. If *S* and *T* are dissimilar, they mostly have short matches, the loci of such short matches have low tree depth in 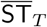, and they do not create long runs in `F`. If *S* and *T* are similar, they have many long matches, the loci of such long matches are more likely to have large tree depth in 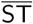, and this can induce long runs in `F`. If the edges of 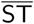 have short labels, there is little to be gained in having *S* similar to *T*. In practice, for pairs of genomes from different species, representing `F` as a delta coding of its runs produces bigger files than just delta-coding every entry of `F` in isolation; but for pairs of human individuals, the run-length encoded `F` is about 15 times smaller (data not shown for brevity). Note that the values in the run-length encoded `F` can be accessed easily using the `runs` bitvector.

We conclude this section by mentioning one more array that is easy to compress. Let `P`_*S,T*_ [1.. |*S*|] be the array that stores at position *i* an arbitrary location of *T* at which *S*[*i*..*i* +`MS`[*i*] − 1] starts^3^ [6]. Rather than storing all locations, it suffices to store just those of the *informative positions*, i.e. of positions *i* such that `MS`[*i*] > `MS`[*i* − 1] −1: for every other position *j*, `P`[*j*] can be reconstructed by recurring on `P`[*j* − 1] + 1. Statistical properties of informative positions in random sequences were described by [32]. In practice, when comparing genomes from different species, approximately half of all positions are informative; this fraction ranges from 0.7 to 0.2 in proteomes, whereas in human individuals it becomes smaller than 0.01 (see Figure 5 in the supplement). One can imagine other position arrays that might be useful in genome comparison, for example an array that stores zero if a match occurs in distinct chromosomes of the text, or the identifier of the only chromosome that contains the match otherwise.

## 5 Querying the MS bitvector

As mentioned, it is natural to formulate questions on the similarity between a substring of the query and the whole text in terms of matching statistics, and this approach has already been used in bioinformatics for detecting horizontal gene transfer and other structural variations between two genomes. In this section we focus on two types of range query, which we implement on the `ms` bitvector: given an interval [*i*..*j*], we want to return either 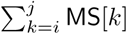 (e.g. to compute a local version of the score by [43]) or max {`MS`[*k*] : *k* ∈ [*i*..*j*]} (e.g. to detect the presence of significant matches).

We answer the max query using the standard approach of dividing the `ms` bitvector into blocks with a fixed number of bits, extracting for each block the maximum MS value that corresponds to a one-bit in the block, and building a range-maximum query (RMQ) data structure on such values^4^. Given a range `MS`[*i*..*j*] in the query, we find the block ′ of ms that contains the *i*-th one-bit, the block *j*′ that contains the *j*-th one-bit, and we query the RMQ data structure on the range of blocks *i*′ + 1..*j*′− 1 (if it is not empty): this returns the index of a block with largest value, thus we perform a linear scan of the returned block, as well as of the suffix of block *i*′ and of the prefix of block *j*′ (if any). We implement this approach using the `rmq_succinct sct` and `bit_vector` data structures from the SDSL library [18]. For ranges *i*..*j* approximately equal to two blocks or larger, this method allows answering a range-max query in the same time as scanning two full blocks and querying the RMQ (see Figure 4); for shorter ranges, it takes the same time as a linear scan of the range. In practice, when the query is the human genome and the block size is, say, 1024 bits, we can answer arbitrary range-max queries in a few milliseconds using just two megabytes for the RMQ and 12 megabytes for the precomputed maximum of each block (if we do not want to compute it on the fly). Other space/time tradeoffs are possible, but we omit a detailed analysis for brevity.

**Figure 4:**
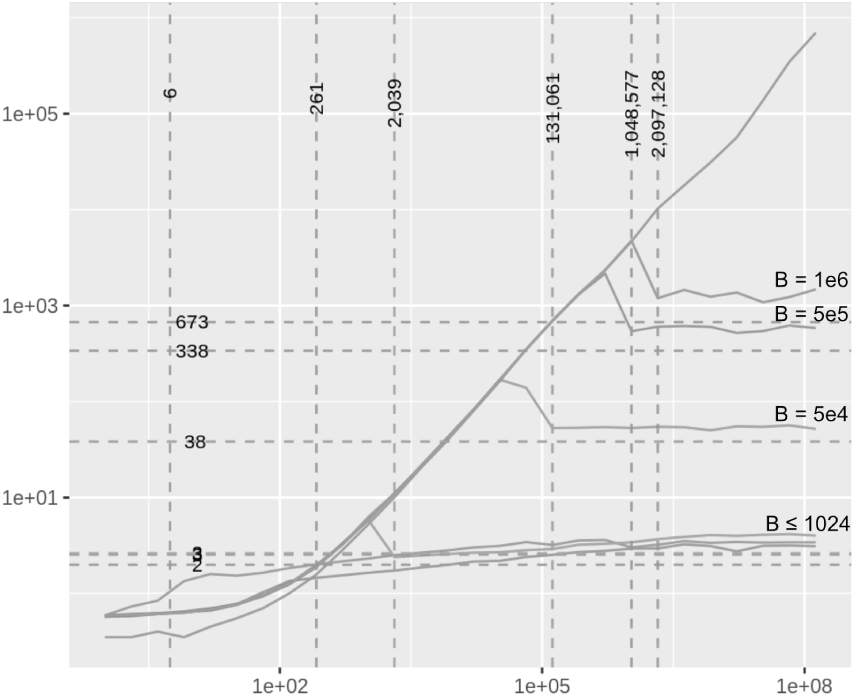
Running time (vertical axis, milliseconds) as a function of range size (horizontal axis) in range-max queries. Diagonal line: baseline, non-optimized scanning of the `bit_vector` data structure from SDSL, without an index. Curves with plateaus: scanning combined with an RMQ index for several settings of block size (indicated by *B*). Every point is the average of 20 random queries of the same size. Vertical dashed lines: query sizes that are approximately equal to two blocks of the `ms` bitvector (the label of a vertical line is the average size in bits of the query range when mapped to the bitvector). Dataset: *Homo sapiens* and *Mus musculus* genomes. Similar trends appear for pairs of genomes from human individuals.

After taking care of some details, this approach can be applied to compressed versions of the `ms` bitvector as well: our implementation supports RRR and run-length encoding (RLE) using the `rrr_vector` and `RLEVector` data structures, from SDSL and from the RLCSA code by [38], respectively. During a scan, issuing one access operation for every bit is clearly suboptimal: instead, in the uncompressed and in the RRR-compressed `ms`, we extract 64 bits at a time and we look up every byte in a precomputed table. This gives speedups between 2 and 8, depending on dataset and range size (see Figure 20 in the supplement). In the RLEcompressed `ms` we process one run at a time, and in very similar strings scanning becomes from a hundred to a thousand times faster than accessing every bit, or more. Overall, scanning the RLE-compressed bitvector of very similar strings processing one run at a time, is approximately *ten times faster than scanning the corresponding uncompressed bitvector* processing 64 bits at a time (see Figure 21 in the supplement). Thus, if the target application is not interested in MS values below some threshold, one might swap the uncompressed bitvector with one of the permuted and RLE-compressed variants described in Section 4, and this might speed up range-max queries at no cost.

As customary, by recurring on the output of RMQ queries one can report all blocks in the range with maximum MS value, or all blocks with MS value at least *τ*, in linear time in the size of the output. Setting the block size to log |*S*| gives an RMQ data structure of *O*(|*S*|*/* log |*S*|) bits, and it allows replacing the linear scan of a block with a constant-time lookup from a table of *o*(|*S*|) bits in which we store the relative location of a largest value inside each block. We use the RMQ to detect all blocks in the range that contain at least one large value, and we use lookups from another table of *o*(|*S*|) bits (which stores offsets between one-bits with MS value at least *τ*) to report all locations with MS value at least *τ* in linear time on their number.

The optimized scanning can be applied to range-sum queries as well, with similar speedups (see Figure 21 in the supplement). To implement a range-sum query [*i*..*j*] over an arbitrary range, we just store the prefix sums that correspond to the last one-bit in every block, and we scan the two blocks that contain the one-bit that corresponds to *i* and the one-bit that corresponds to *j*. Scanning can also be used to implement other primitives in analytics, like plotting all MS values in a range or their histogram, computing the position *k* of a longest interval [*k*..*k* + `MS` [*k*] −1] that contains [*i*..*j*], or finding all windows of fixed length *k* inside [*i*..*j*] with maximum sum.

## Acknowledgements

We thank Giorgio Vinciguerra for help with the software by [5], and Massimiliano Rossi and Dominik Koeppl for help with the software by [6].

## Footnotes

1 Actually, since at iteration j we compute up to 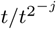 total solutions, we could afford to allocate 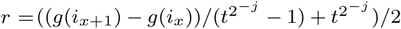 processors per solution.

2 In some applications *τ* might even change along *S*, e.g. when *S* is the concatenation of several genomes with different similarity to *T*.

3 One could of course define lossy variants in which the locations of short matches are discarded.

4 Clearly the maximum of each block is assumed by an informative position defined in Section 4.1, thus it suffices to consider just those during construction.

