## Supplementary material for "Fast and compact matching statistics analytics"

### 1 Details on the experiments

We run our parallelism experiments on a server with four Intel Xeon E7-4830v3 processors, each with 12 cores, a base frequency of 2.1 GHz, and a maximum frequency of 2.7 GHz. The machine has one terabyte of RAM and no other user. We compile our code with `gcc 6.2.0`.

Since our parallel algorithm might not scale well when the query is very similar to the text, we experiment with the following pairs of similar genomes and proteomes: *Arabidopsis lyrata* and *Arabidopsis thaliana*; *Zea mays* and *Oryza sativa*; *Rattus norvegicus* and *Mus musculus*; *Gallus gallus* and *Taeniopygia guttata*; *Brugia malayi* and *Caenorhabditis elegans*; *Drosophila melanogaster* and *Anopheles gambiae*; *Pan troglodytes* and *Homo sapiens*. When building each dataset, we download the latest assembly from NCBI and we concatenate its sequences using a separator that is not in the alphabet, replacing runs of undetermined characters with a single occurrence of the separator. To probe the high-similarity regime, we compare 28 random pairs of human chromosome 1 taken from the 1000 Genomes Project[1]. We transform the set of variants from each individual into two full copies of chromosome 1, of length approximately 215 million each, using the `vcf2multialign` tool[2].

We run our sequential range query experiments on a server with two Intel Xeon E5-1650v4 processors, each with 6 cores, a base frequency of 3.6 GHz, and a maximum frequency of 4 GHz. The machine has 128 GB of RAM and no other user. We compile our code with `gcc 8.4.0`.

In some experiments we need to approximate a use case in which the genomes or proteomes of different species are compared to one another: for this we use a dataset of 24 eukaryotic species with relatively large genomes that has been used for whole-genome phylogeny reconstruction by average common substring before [7]. Such genome files range from approximately 87 million to 5.8 billion characters. We call this the *ACS dataset* in what follows.

[1] `ftp://ftp.1000genomes.ebi.ac.uk/vol1/ftp/release/20130502/ALL.chr1.phase3_shapeit2_mvncall_integrated_v5a.20130502.genotypes.vcf.gz`

[2] `https://github.com/tsnorri/vcf2multialign`

### 2 Supplemental figures

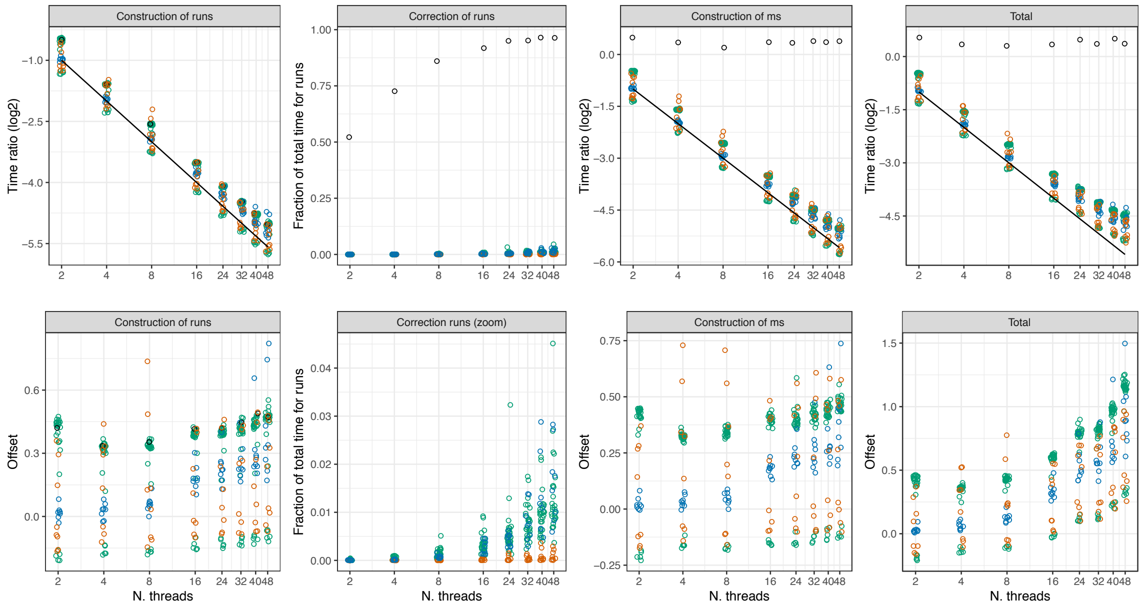


Figure 1: Running time of the parallel implementation as the number of threads increases. Red: genomes of similar species; blue: proteomes of similar species; green: pairs of human chromosome 1 from distinct random individuals; black circles: identical query and text (human chromosome 1 from two random individuals); black line: $1/t$, where $t$ is the number of threads. Time ratio: time of the parallel implementation divided by the time of the sequential implementation. Offset: $(r - 1/t)/(1/t)$, where $r$ is the time ratio and $t$ is the number of threads. Correction of `ms` is not shown since it is negligible in all cases.

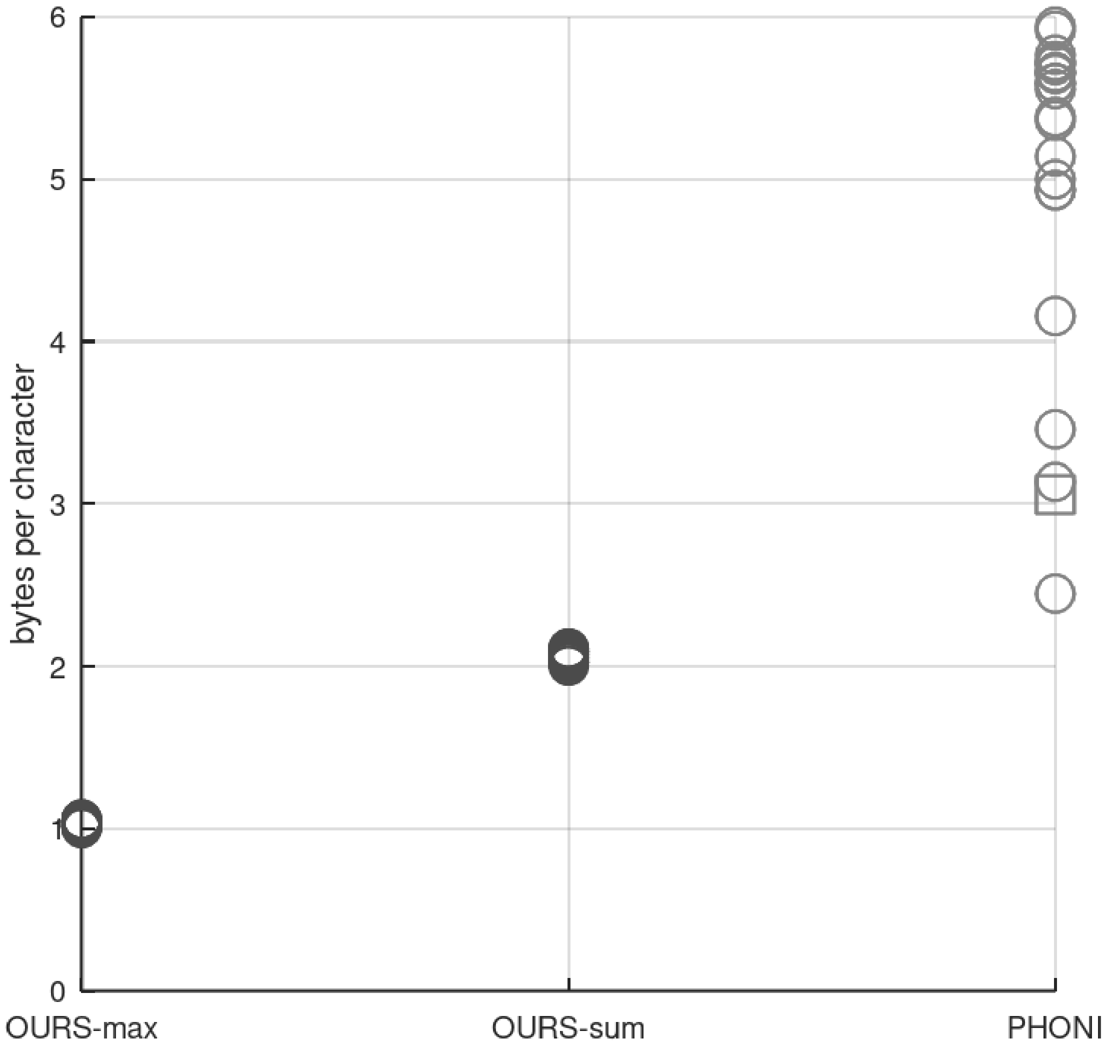


Figure 2: Comparing the size of the indexes used by our implementation (“OURS”, [1]) to those used by the implementation in [3] (“PHONI”), on all genomes in the ACS dataset. Since our algorithm needs just the index of one direction at any time, we report such value in “max”, whereas “sum” is the size of all our indexes in both directions. The square is the size of the index in [3] built on the concatenation of all the approximately 8600 bacterial genomes in NCBI (34 billion characters): such a string should be moderately repetitive.

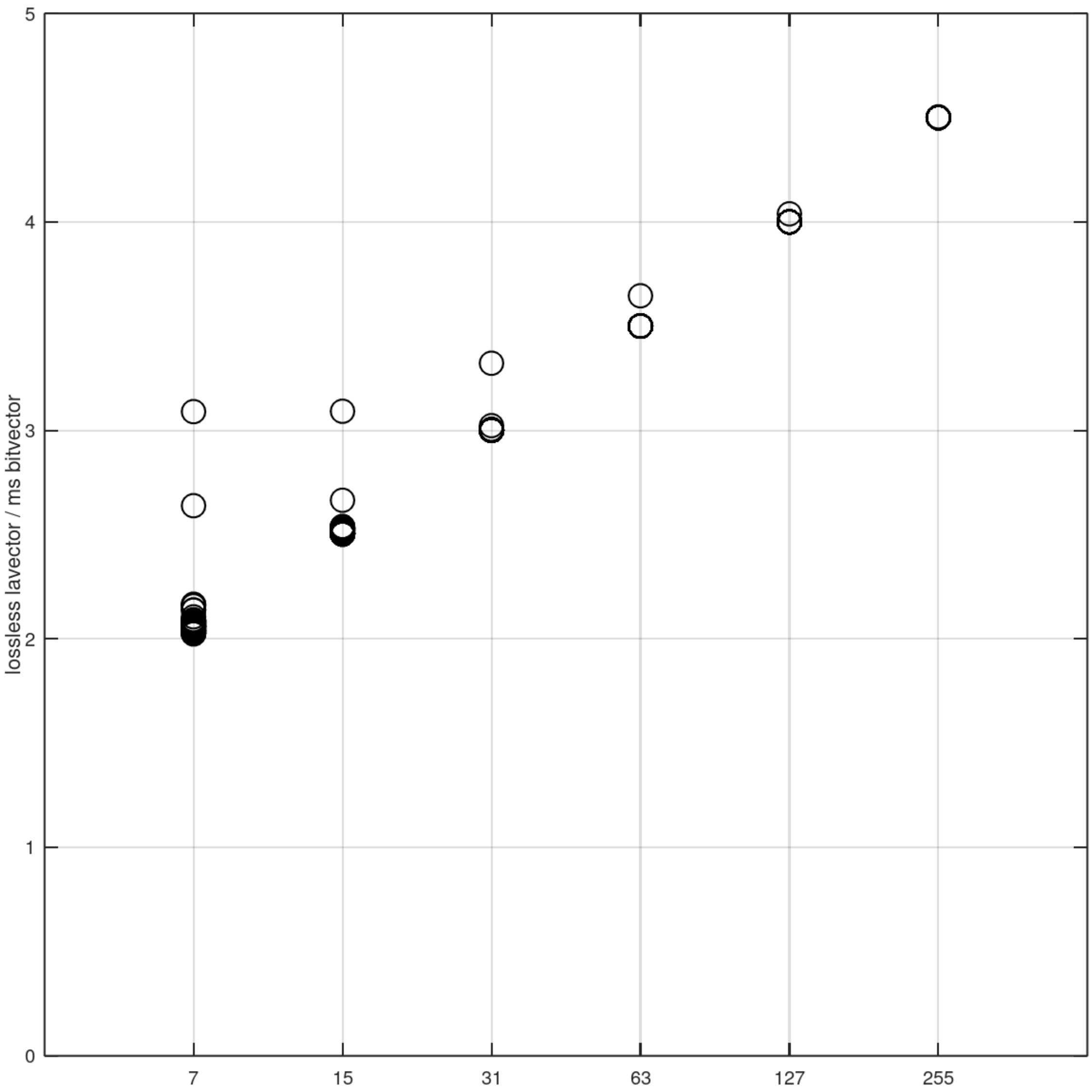


Figure 3: Size of the lossless `la_vector` implementation in [2] applied to the `ms` bitvector of some pairs of large genomes in the ACS dataset, compared to the original size of the `ms` bitvector. Horizontal axis: the maximum error $(2^{c-1}-1)$ that can be represented by a correction encoded in $c$ bits.

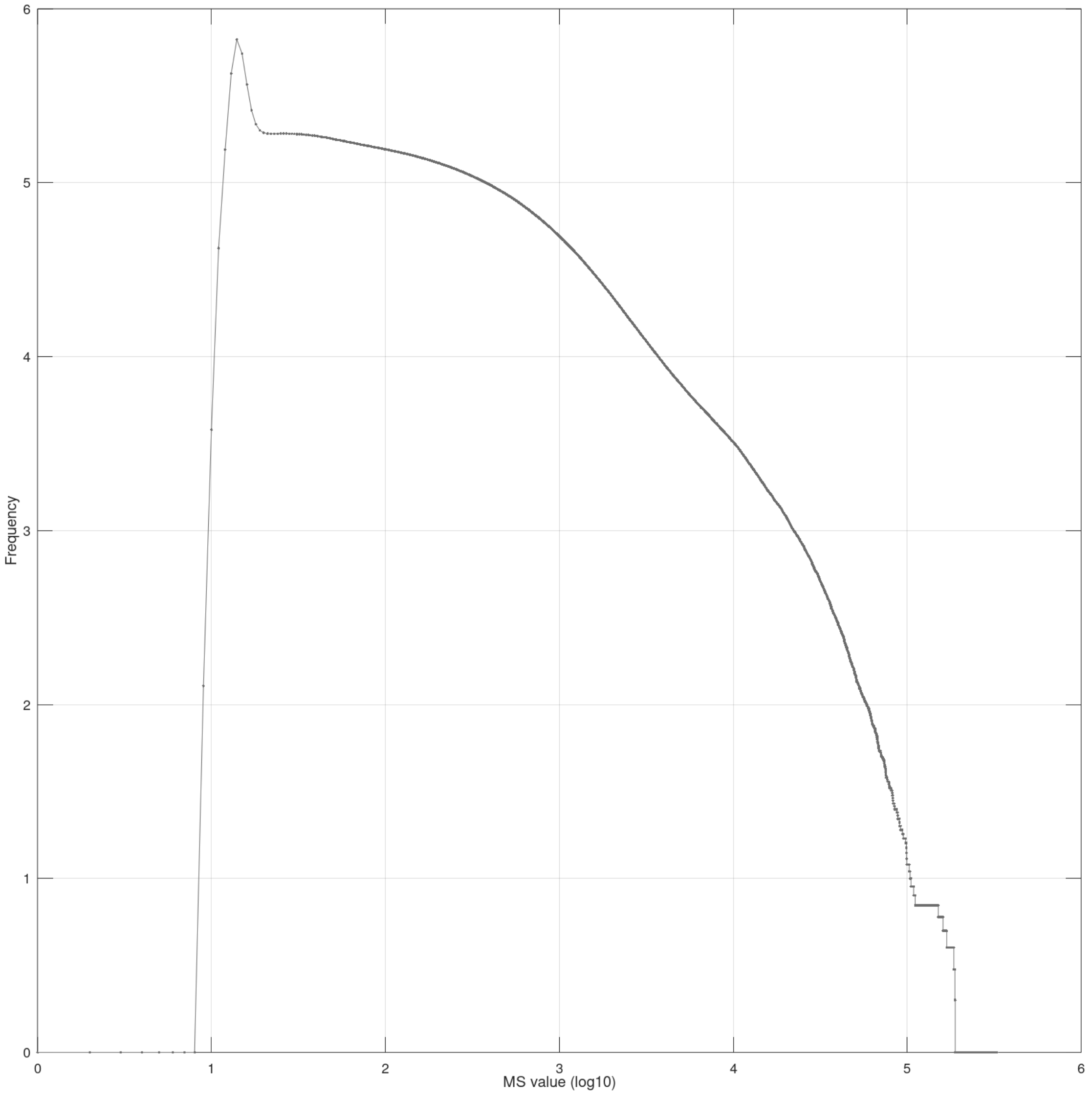


Figure 4: Histogram of MS values between *Homo sapiens* and *Pan troglodytes*. The vertical axis is in logarithmic scale (base 10). The histogram for pairs of unrelated genomes is similar up to the peak, but thereafter it goes quickly to zero instead of having a long tail.

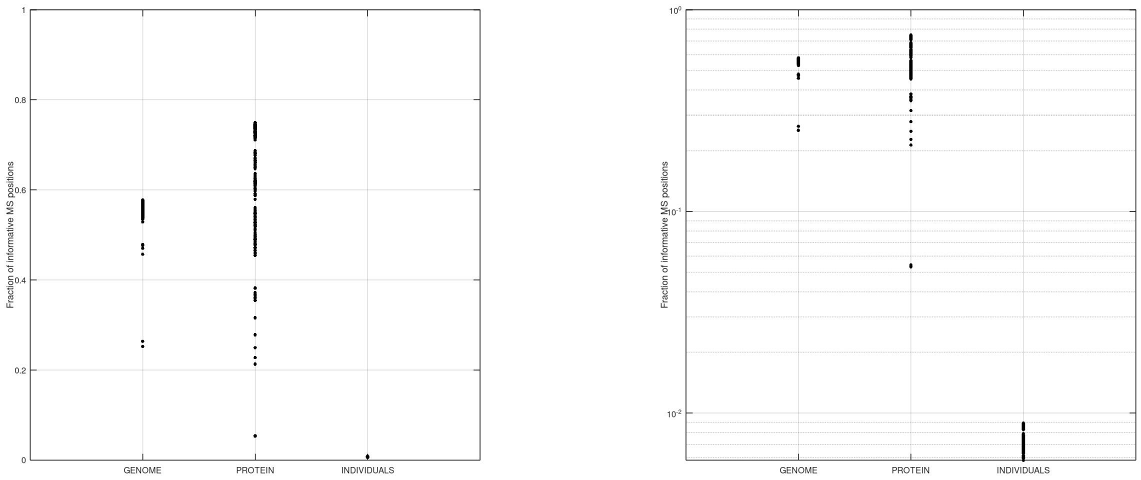


Figure 5: Fraction of *informative positions*, i.e. of positions $i$ such that $\mathsf{MS}[i] > \mathsf{MS}[i-1]-1$, for every pair of sequences in the ACS dataset ("genome" and "protein") and in the dataset of human individuals. The right panel shows the same data as the left one, but in logarithmic scale.

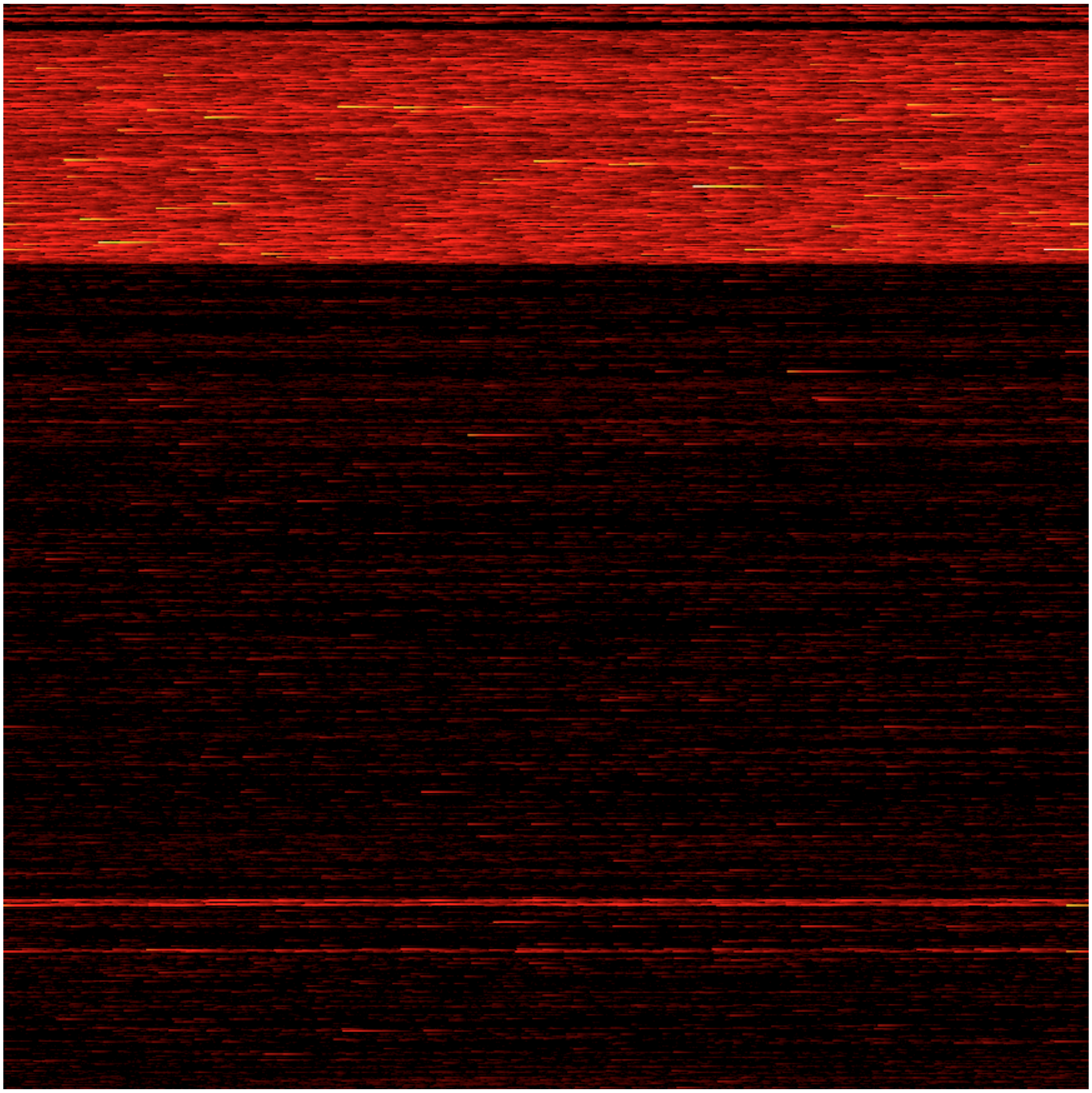

Figure 6: The matching statistics array highlights large-scale patterns of similarity. The figure shows the first hundred million values of the MS array between *Homo sapiens* (query) and *Pan troglodytes* (text). The array is displayed in several rows of fixed length just for visualization purposes. Every pixel is a position in MS. Colors represent MS values (black < red < yellow < white), which range between 5 and 1442.

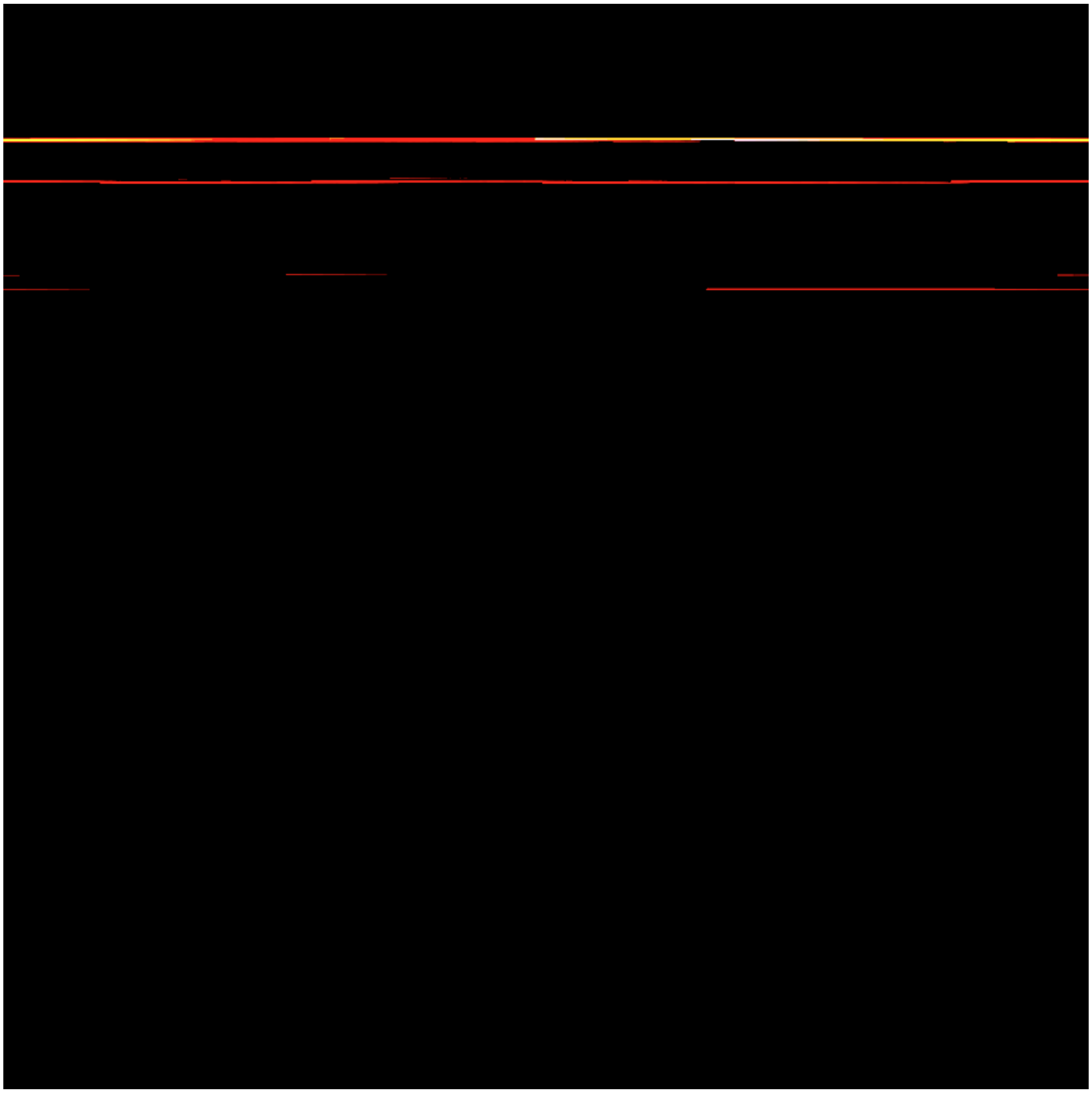

Figure 7: Same as Figure 6, but for *Homo sapiens* (query) and the reverse-complemented of itself (text). MS values range between 5 and 12.4k.

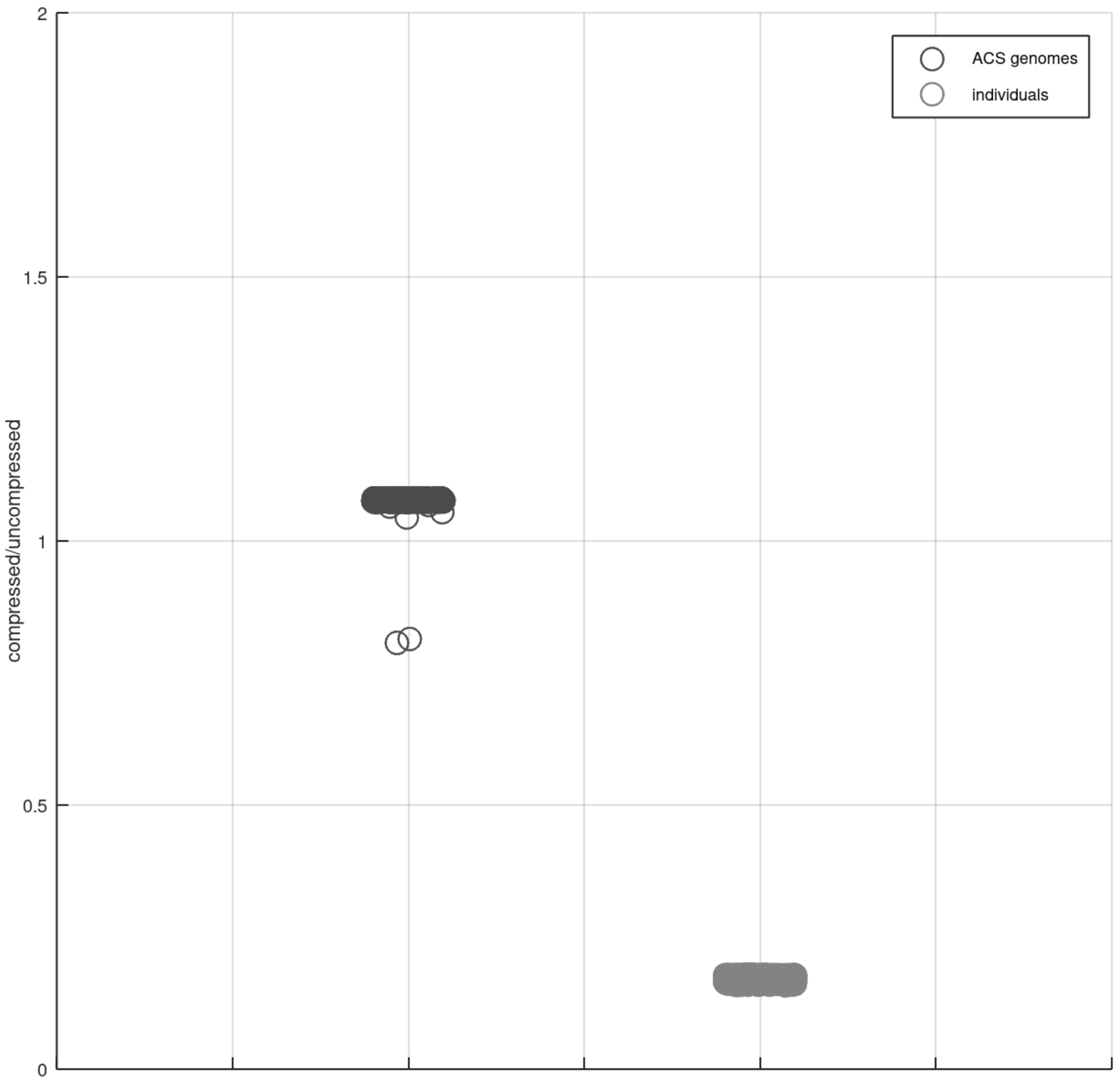


Figure 8: Compressing the `ms` bitvectors of genomes using the `rrr_vector` data structure from the SDSL library [4]: ACS dataset (left) and human individuals dataset (right). Most bitvectors of different species get expanded.

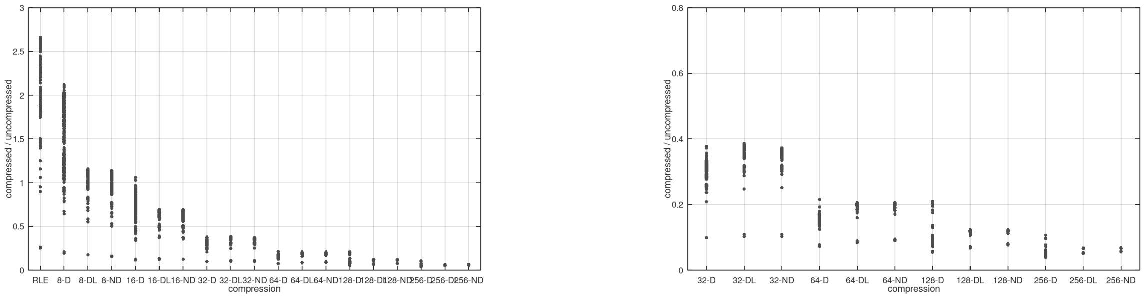


Figure 9: Same as Figure 3 in the paper, but for pairs of proteomes from different species in the ACS dataset. Negative values of MS are not allowed in the lossy variants. The right panel is a zoom-in of the left panel.

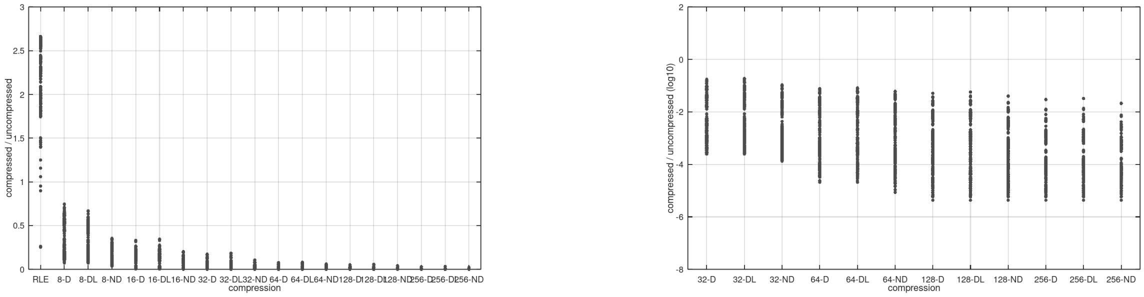


Figure 10: Same as Figure 9, but in this case negative values of MS are allowed.

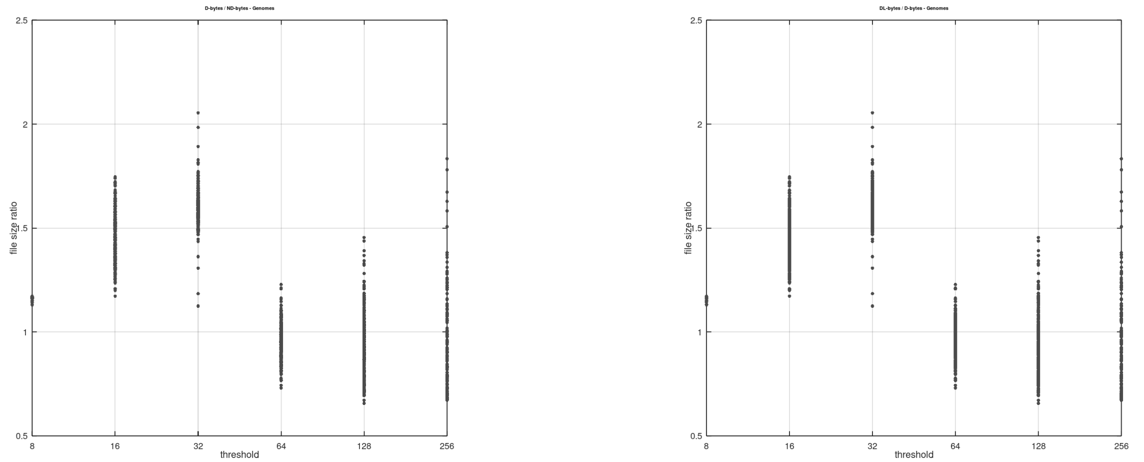


Figure 11: Ratios of D over ND (left panel) and of DL over D (right panel), for every pair of genomes from different species in the ACS dataset. Negative MS values are not allowed. Sizes are measured on disk, and refer to the `RLEVector` data structure by [6].

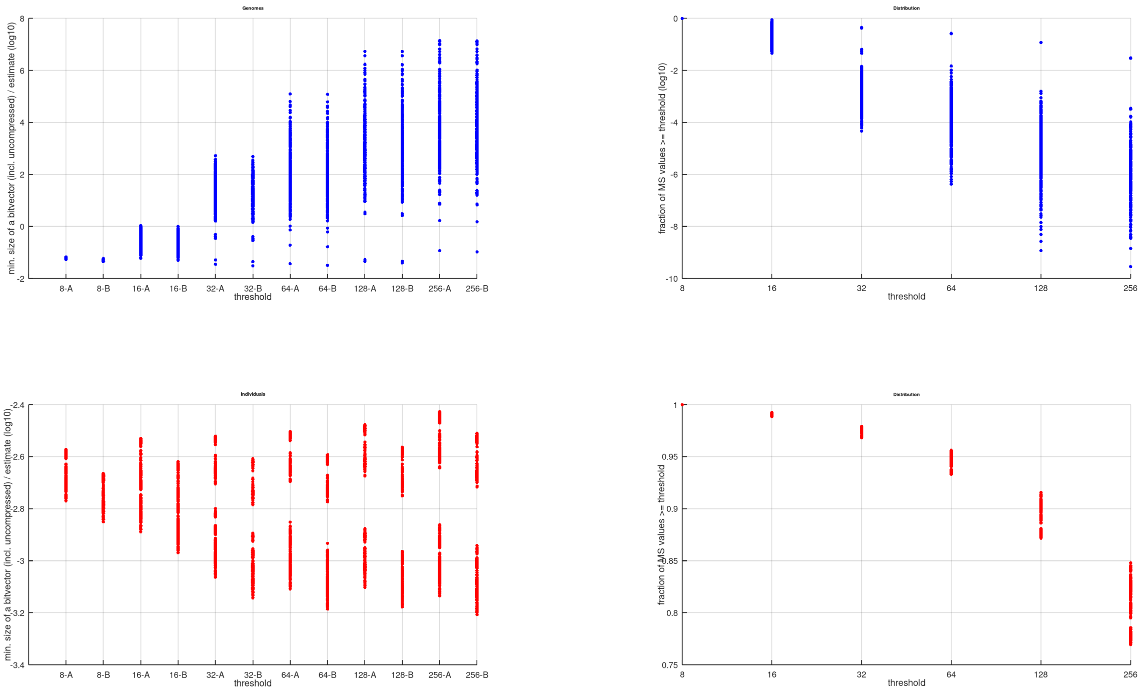


Figure 12: Comparing our lossy schemes to a simple baseline that stores just the MS values above threshold (see Section 4 in the paper for details). Top panels: genomes from different species in the ACS dataset. Bottom panels: genomes from the human individuals dataset. Right panels: fraction of values above threshold. Left panels: ratio between the smallest disk size of one of our bitvectors (i.e. the minimum over D, DL, ND, and not permuted), and the disk size of the $A_\tau$ and $B_\tau$ schemes.

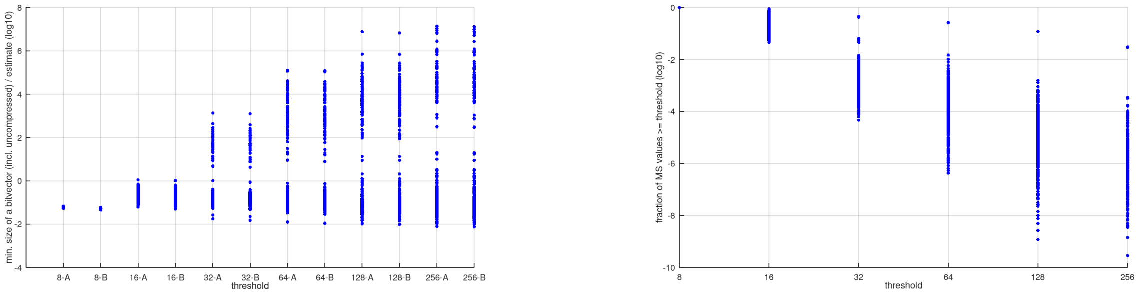


Figure 13: Same as Figure 12 (top), but allowing negative values in the permuted bitvectors.

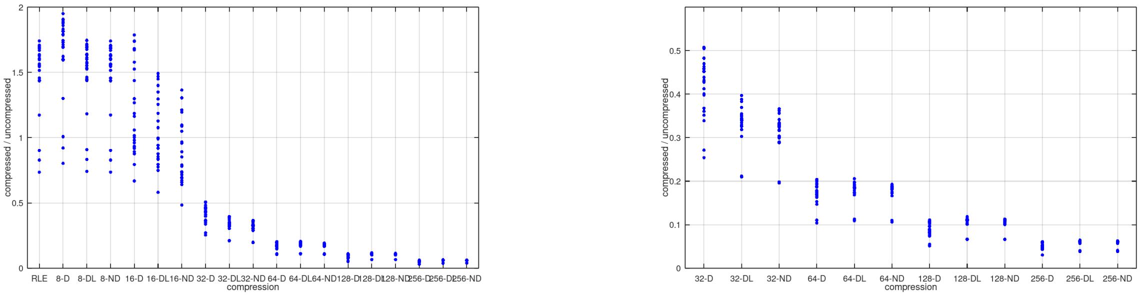


Figure 14: Same as Figure 3 in the paper (left panel), but for the PLCP array of the genomes in the ACS dataset. The right panel is a zoom-in of the left panel.

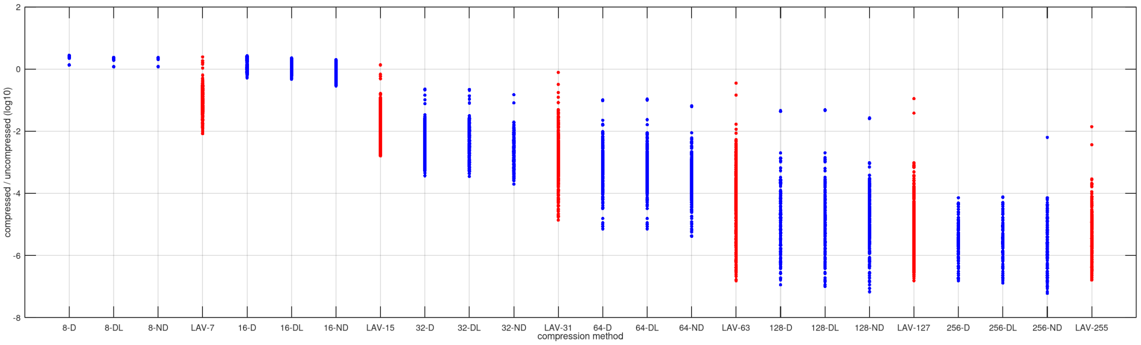


Figure 15: Comparing our lossy schemes to a lossy variant of `la_vector` on the ACS genome dataset. $\tau$-D, $\tau$-DL, $\tau$-ND: disk size of our bitvector permutations with threshold $\tau$, negative values allowed, and encoded with the `RLEVector` data structure by [6] (blue). LAV-$\epsilon$: disk size of a lossy variant of the `la_vector` implementation [2] in which correction values are not stored (red). $\epsilon$ is the maximum error ($2^{c-1}-1$) that can be introduced when discarding the corrections (encoded in $c$ bits).

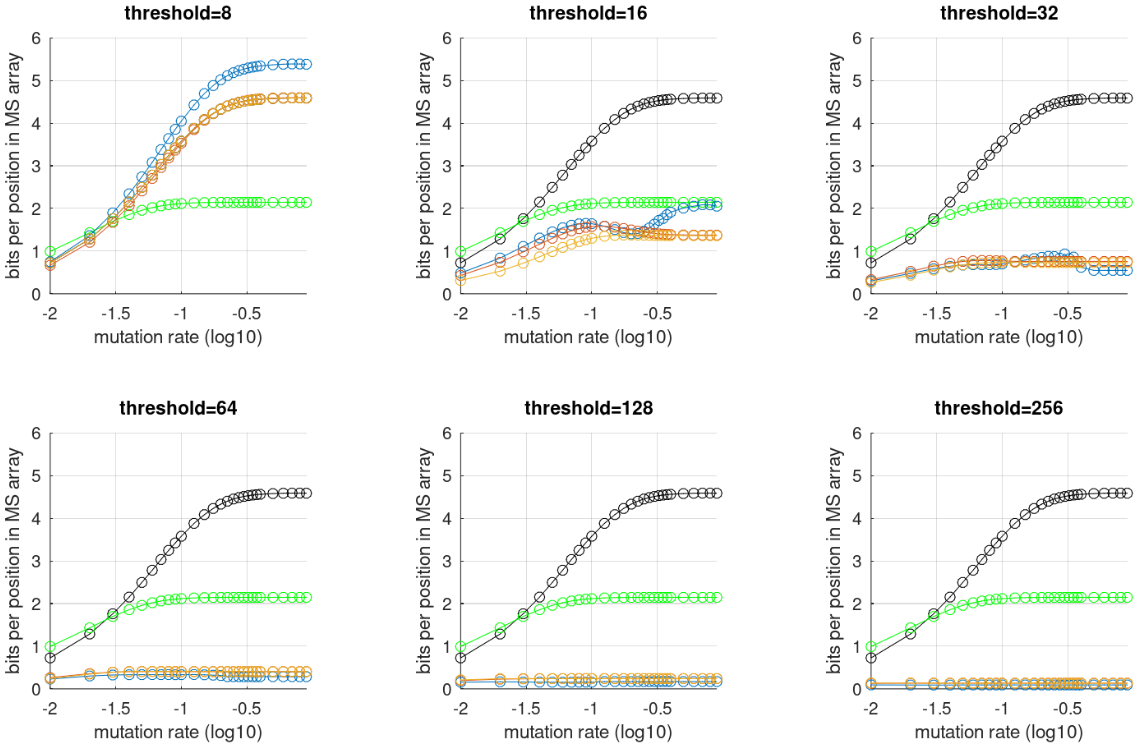


Figure 16: Efficiency of our lossy compression schemes with varying mutation rate between query and text. Text: human chromosome 1. Query: a 10 Mbps prefix of the text, with added mismatches at variable rate (horizontal axis). Green: disk size of the `rrr_vector` data structure from the SDSL library [4] built on the original `ms` bitvector. Black: disk size of the `RLEVector` data structure by [6] built on the original `ms` bitvector. Other colors: disk size of `RLEVector` built on the D (blue), DL (red) and ND (yellow) permutations. Negative values of MS are not allowed. The uncompressed `ms` bitvector takes 2 bits per position, so plain RLE *expands* the bitvector for mutation rates $\geq 0.04$.

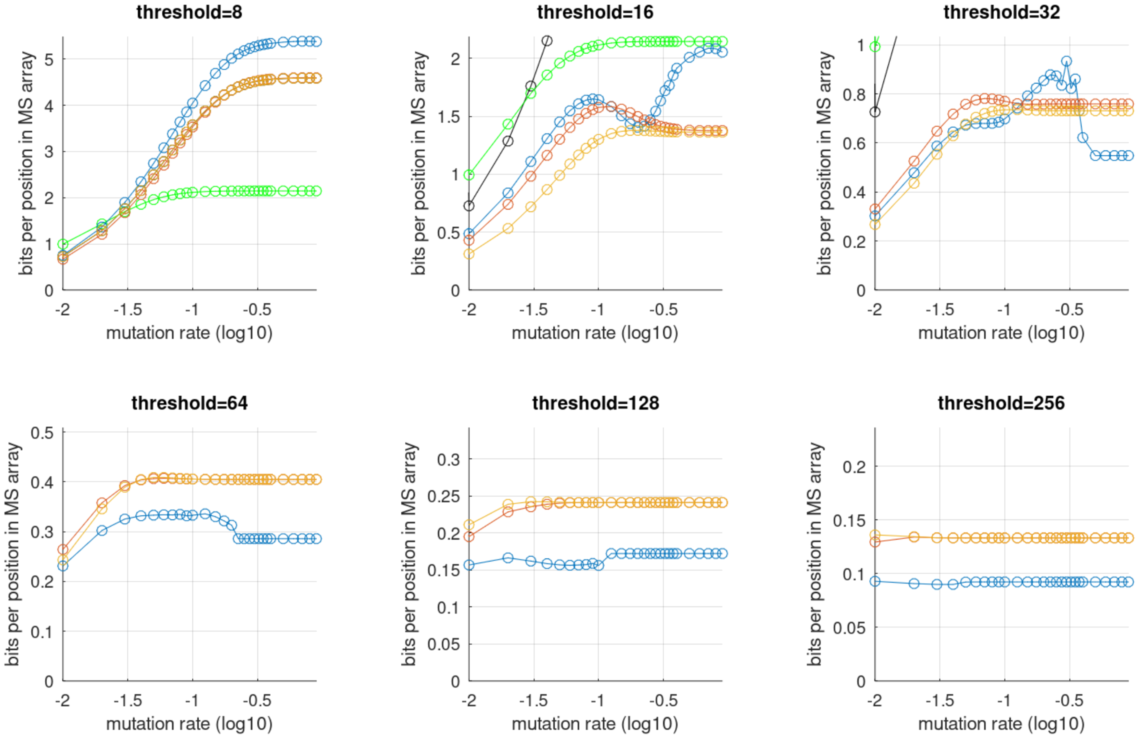


Figure 17: Zoom of Figure 16. At high mutation rates (dissimilar sequences), DL and ND (red and yellow) are almost equivalent, and D (blue) becomes useful only for high values of the threshold. At low mutation rates (similar sequences), ND is equivalent to DL or preferable to it for all values of the threshold.

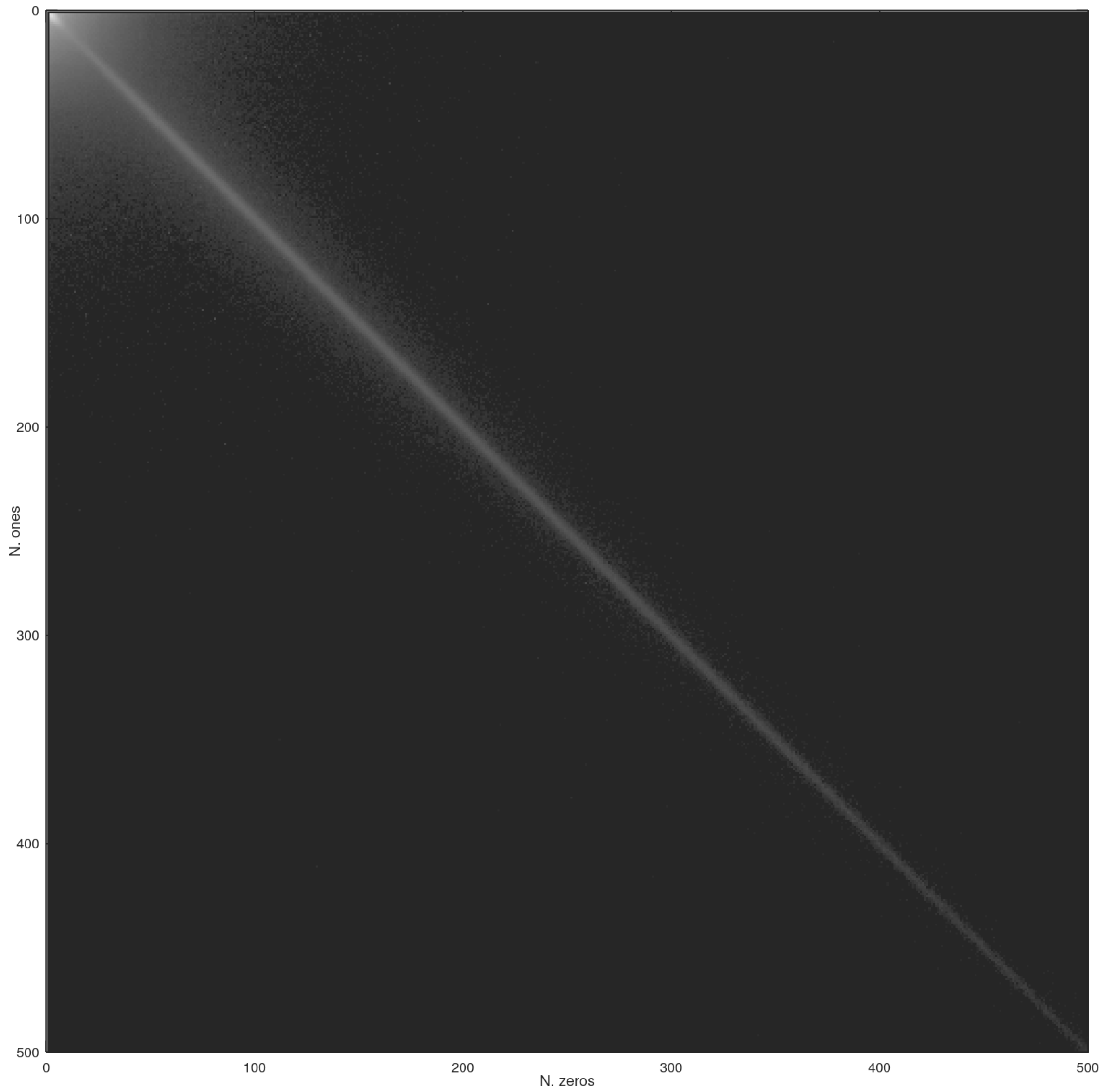


Figure 18: Correlation between the length of a run of zeros and the length of the following run of ones in the `ms` bitvector of *Homo sapiens* and *Pan troglodytes*. Every pair of lengths is a cell of the matrix; the value in a cell is the logarithm of the number of observed pairs (black means low, white means high). We detected no correlation between the length of a run of zeros and the length of the *preceding* run of ones (or the length of any of the preceding runs of zeros). A similar trend appears in the bitvector encoding of the PLCP array of the human genome (data not shown) [5].

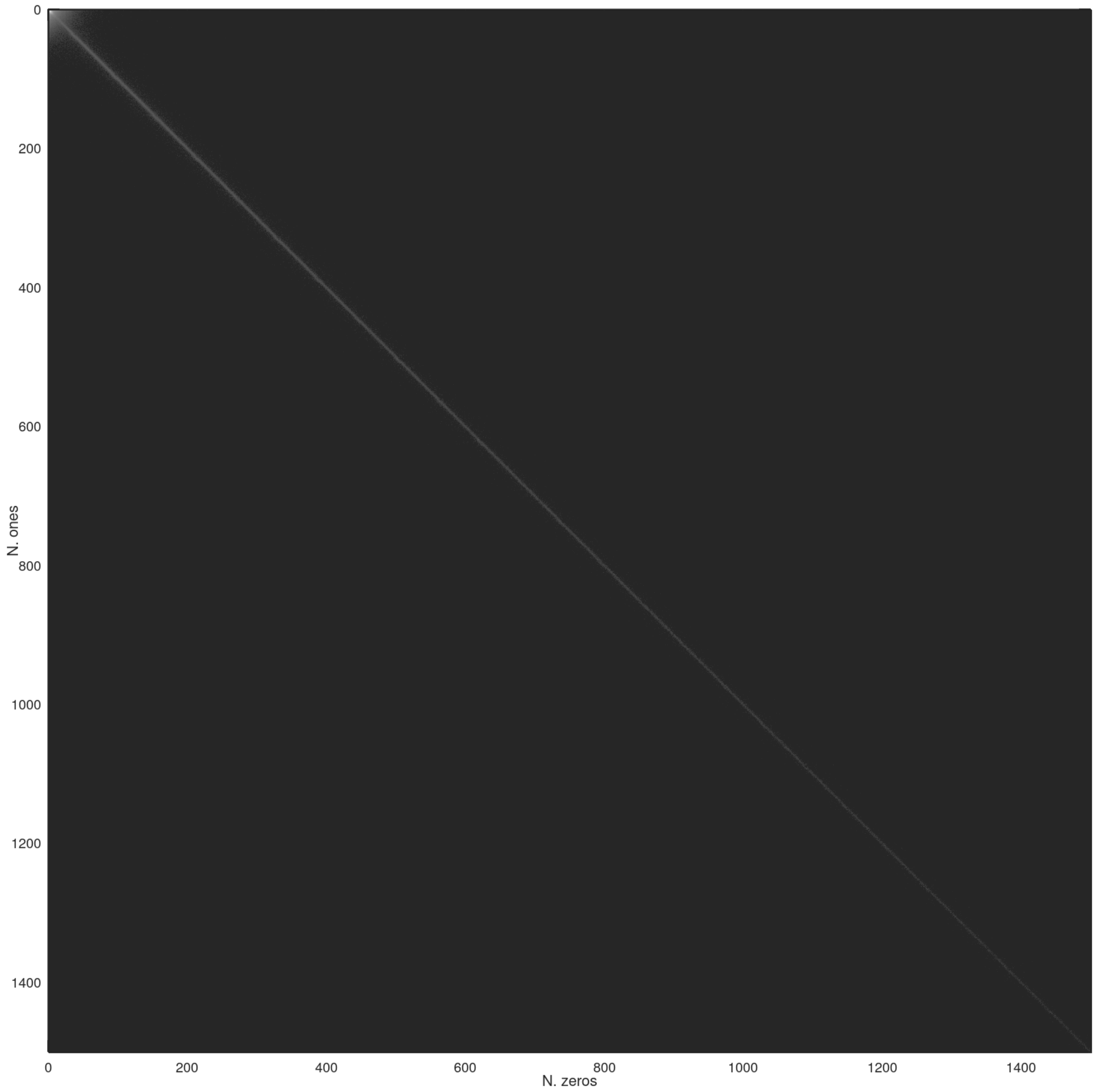


Figure 19: Same as Figure 18, but for a pair of chromosome 1 sequences from human individuals.

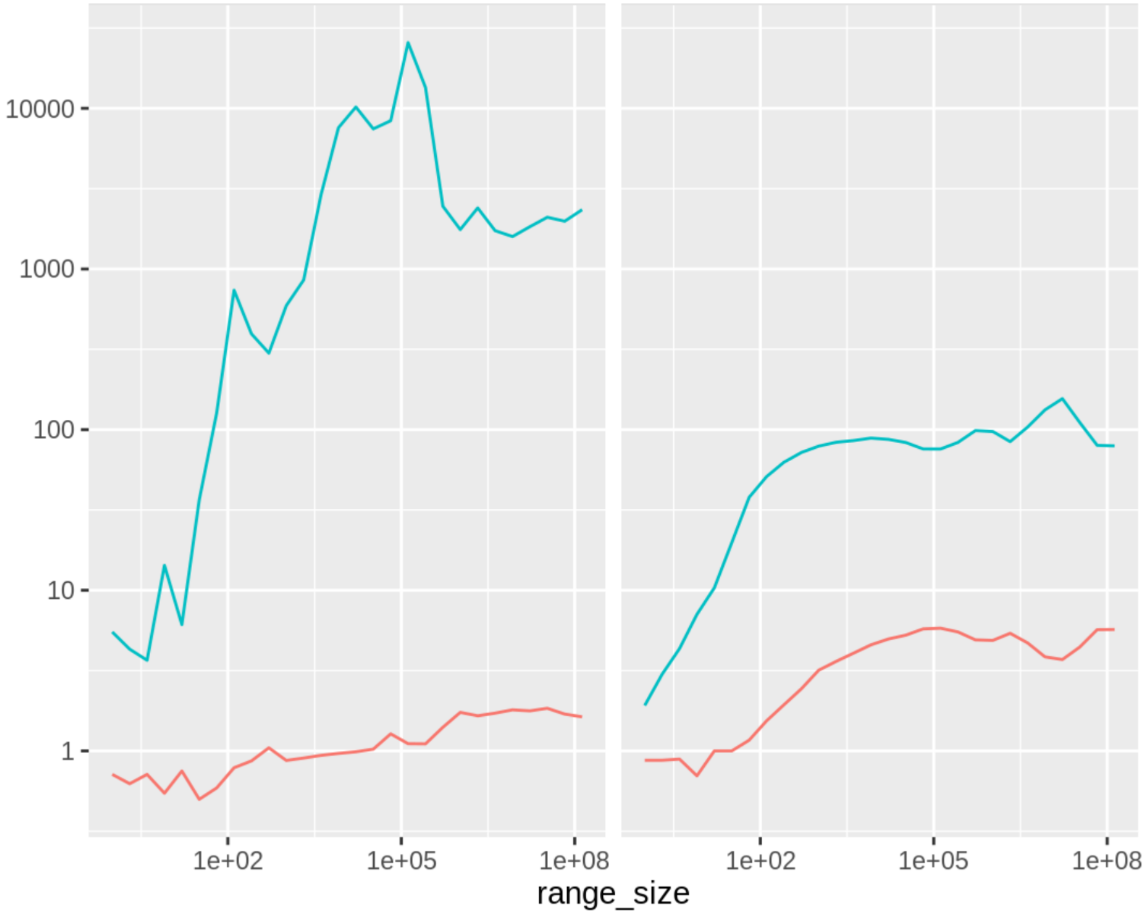


Figure 20: Scanning the `ms` bitvector in range-max queries: ratio between the time of the baseline approach that accesses every bit of the bitvector, and the time of our optimized code. Red: `bit_vector` data structure from SDSL [4]. Green: `RLEVector` data structure from [6]. Left: chromosome 1 of two human individuals. Right: *Homo sapiens* and *Mus musculus*. Every point is the average of 50 random queries of the same size.

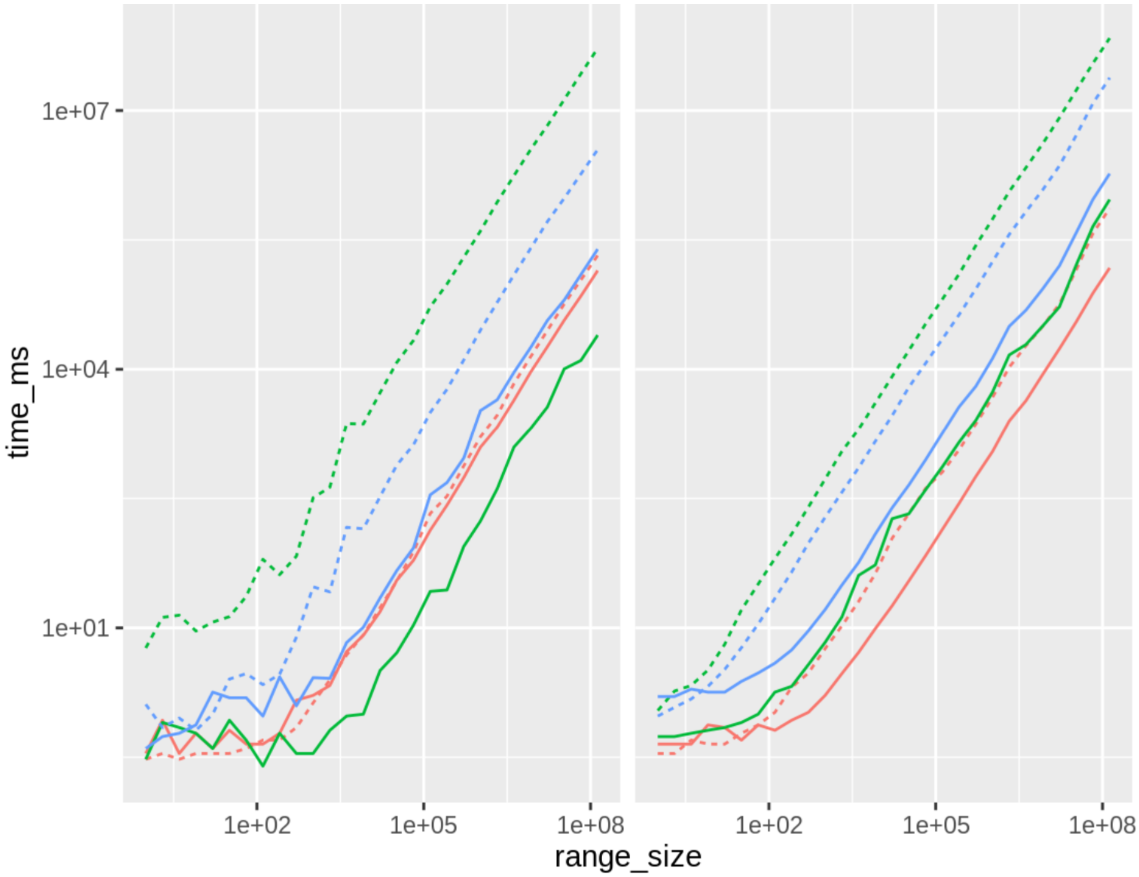


Figure 21: Scanning the `ms` bitvector in range-sum queries: time of the baseline approach that accesses every bit of the bitvector (dotted lines), and time of our optimized code (continuous lines). Red: `bit_vector` data structure from SDSL [4]. Green: `RLEVector` data structure from [6]. Blue: `rrr_vector` from SDSL. Left: chromosome 1 of two human individuals. Right: *Homo sapiens* and *Mus musculus*. Every point is the average of 50 random queries of the same size.

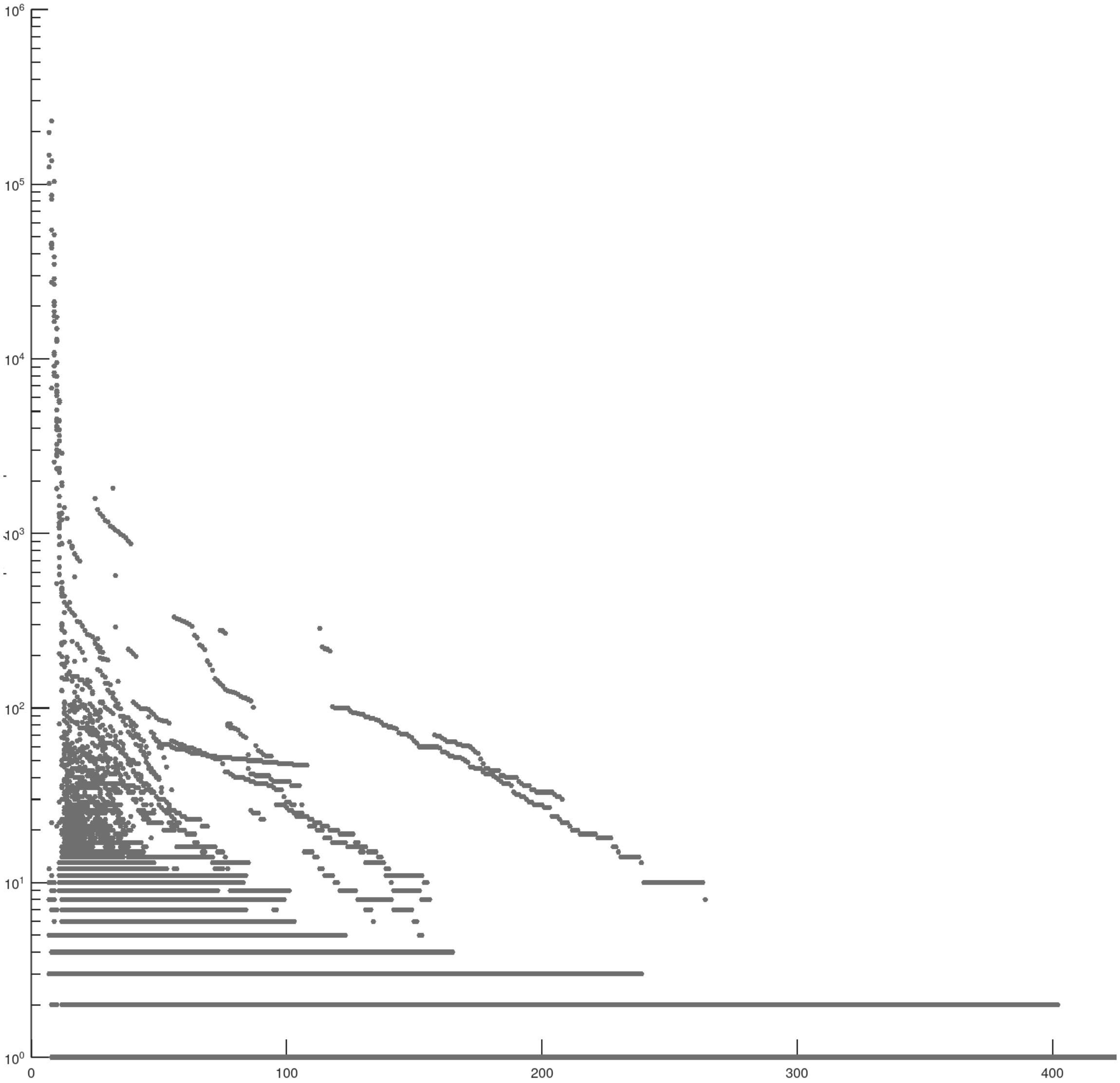


Figure 22: Long matching statistics values between the query (*Homo sapiens*) and the text (*Pan troglodytes*) can have high frequency in the text. Horizontal axis: matching statistic value. Vertical axis: frequency in the text (logarithmic scale). This suggests that the two genomes share exact repeats.

### References


[1] Djamal Belazzougui, Fabio Cunial, and Olgert Denas. Fast matching statistics in small space. In *Proceedings of the 17th International Symposium on Experimental Algorithms (SEA 2018)*, volume 103. Schloss Dagstuhl-Leibniz-Zentrum fuer Informatik, 2018.
